# Functional genomics and tumor microenvironment analysis reveal prognostic biological subtypes in Mantle cell lymphoma

**DOI:** 10.1101/2025.06.25.661613

**Authors:** Sunandini Sharma, Roshia Ali, Alyssa Bouska, Dylan Jochum, Meghana Kesireddy, Simeon Mahov, Joseph Lownik, Weiwei Zhang, Waseem Lone, Mahfuza Afroz Soma, Alicia Gamboa, Vaishnavi Devarkonda, Dalia Elgamal, Atqiya Fariha, Adnan Mansoor, Douglas Stewart, Peter Martin, Brian K. Link, Ranjana H. Advani, Paul M. Barr, Andre H. Goy, Amitkumar Mehta, Manali Kamdar, Deborah M. Stephens, Veronika Bachanova, Lynette Smith, Ryan Morin, Prasath Pararajalingam, Matthew A. Lunning, Kai Fu, Dennis Wiesenberger, Wing C Chan, Joseph Khoury, Timothy C. Greiner, Julie M Vose, Akil Merchant, Chengfeng Bi, Javeed Iqbal, the North American Mantle Cell Lymphoma Project (NAMCLP)

**Author notes:** Correspondence: Chengfeng Bi, MD, PhD, Division of Oncology & Hematology, Department of Internal Medicine, University of Nebraska Medical Center, Omaha, NE 68198;., Javeed Iqbal, PhD, MS, Department of Pathology and Microbiology, and Immunology, University of Nebraska Medical Center, Omaha, NE 68198-6842,.

## Abstract

Mantle cell lymphoma (MCL) is a genetically and clinically heterogeneous B-cell malignancy. We studied two MCL cohorts with differing treatment patterns: one enriched for immunochemotherapy, the other for chemotherapy alone. *TP53* alterations were consistently associated with poor prognosis, whereas *ATM* mutations correlated with improved outcomes following rituximab-based chemotherapy. Based on recurrent genetic events, six clusters were identified and refined into three prognostic groups: high-risk (*TP53* mutations and deletions at 17p13.3, 13q14.2, and 19p13.3), intermediate-risk (*ATM* and epigenetic regulator mutations, or gains at 8q/17q/15q), and low-risk (lacking *TP53* alterations, rare *ATM* mutations without 11q deletions, gains at 3q, deletions at 6q). Transcriptomic analysis revealed enrichment of proliferation, metabolism-promoting gene signatures in high-risk; angiogenesis and NOTCH signaling in intermediate-risk; and proinflammatory-related (i.e., IFNα, TNFα) in low-risk MCLs. Multi-proteomic spatial profiling using imaging mass cytometry (IMC) demonstrated enrichment of CD8⁺ T cells with high expression of exhaustion markers and dominant population of myeloid cells skewed toward an M2-like phenotype. Compared to *ATM*-perturbed tumors, *TP53*-perturbed tumors exhibited enriched SOX11⁺ tumor cells and enhanced tumor-immune cell interactions. Functional analysis revealed that p53 represses BCR signaling through *PTPN6* activation. Collectively, these findings highlight distinct molecular and immune landscapes and reveal therapeutic vulnerabilities in high-risk *TP53*-altered MCL.

## Introduction

MCL is an incurable non-Hodgkin lymphoma (NHL), characterized by the *t*(11;14) involving *IgH::CCND1* rearrangement leading to G_1_-cyclin (CCND1) upregulation and is often associated with SOX11 overexpression^1–3^. While immune-chemotherapeutic regimens have improved clinical outcomes, relapses are frequent, and most patients eventually succumb to the disease^4,5^. The clinical course is highly variable, underscoring the necessity for an improved understanding of disease biology to improve clinical management. Beyond clinicopathological risk stratification, subclassification at the molecular level may provide improved biological and therapeutic justification. Gene expression profiling (GEP) studies identified transcriptional characteristics associated with the aggressive behavior of tumor cells, and established mRNA and miRNA signatures to differentiate MCLs with varying clinical outcomes^6–9^. DNA copy number variation (CNV) analysis has identified virtually exclusive 17p13.3 and 11q22.3 aberrations, highlighting the recurrent deletions of *TP53* and *ATM*^10^. Epigenomic studies revealed MCL subtypes with distinct lineage origin and clinical outcomes^11^. Genetic studies have highlighted DNA damage repair (*ATM*), cell cycle (*TP53, POT1, ASPM*), and chromatin modifiers (*SMARCA4, KMT2D, KMT2C*) as major oncogenic events^12,13^. These findings suggest that *CCND1* overexpression may exert selective pressure on pre-GC-B cells to inactivate the DNA-damage response (DDR) pathway (i.e., *ATM or TP53*) or epigenetic dysregulation, but the underlying biological or clinical heterogeneity remains unexplored.

Although the role of the tumor microenvironment (TME) in MCL pathogenesis remains insufficiently characterized, emerging studies have highlighted a strong interplay between stromal cells and malignant B cells, leading to chronic activation of the BCR-PI3K-mTOR signaling pathway and conferring a survival advantage to the tumor^14,15^. We assembled two MCL cohorts: one rituximab-limited and the other rituximab-enriched (n=153), with the latter treated under intensified protocols (i.e., R-CHOP/R-DHAP or R-Hyper-CVAD/MA)^16^. We aimed to explore MCL pathogenesis using multi-omics approaches, including high-throughput genomics, transcriptomics, and multi-proteomics IMC, and defined the intrinsic genetic alterations of the tumor that determine clinical outcome, in association with the TME. Mechanistically, we investigated the functional role of *TP53* genetic alterations, with the goal of improving disease management in this subtype.

## Results

### Patient cohort and clinical characteristics

The study included two cohorts: Cohort 1 (Coh-1; WES, *n* = 153) and Cohort 2 (Coh-2; Targeted Sequencing [TS], *n* = 137). Clinical and assay details are summarized in **Table 1** and **Supplementary Table 1**, respectively. Rituximab (immunochemotherapy) was part of the primary therapy in 87.1% of Coh-1 and 36.8% of Coh-2 cases. All Kaplan-Meier analyses for Coh-1 excluded Rituximab-naïve cases, and combined cohort outcome analyses included only Rituximab-treated cases. Over 70% of Coh-1 and Coh-2 cases were formalin-fixed, paraffin-embedded (FFPE). For IMC analysis, we included 50 cases: 42 with available sequencing, 5 from the NAMCLP cohort without sequencing, and 3 tissue samples from tonsil, lymph node, and bone marrow. Coh-1 cases included RNA-Seq data from 47 samples, while Coh-2 analysis was based on microarray data (HG-U133plus2 platform) of 25 cases. The study was approved by institutional ethics committees.

**Table 1:**
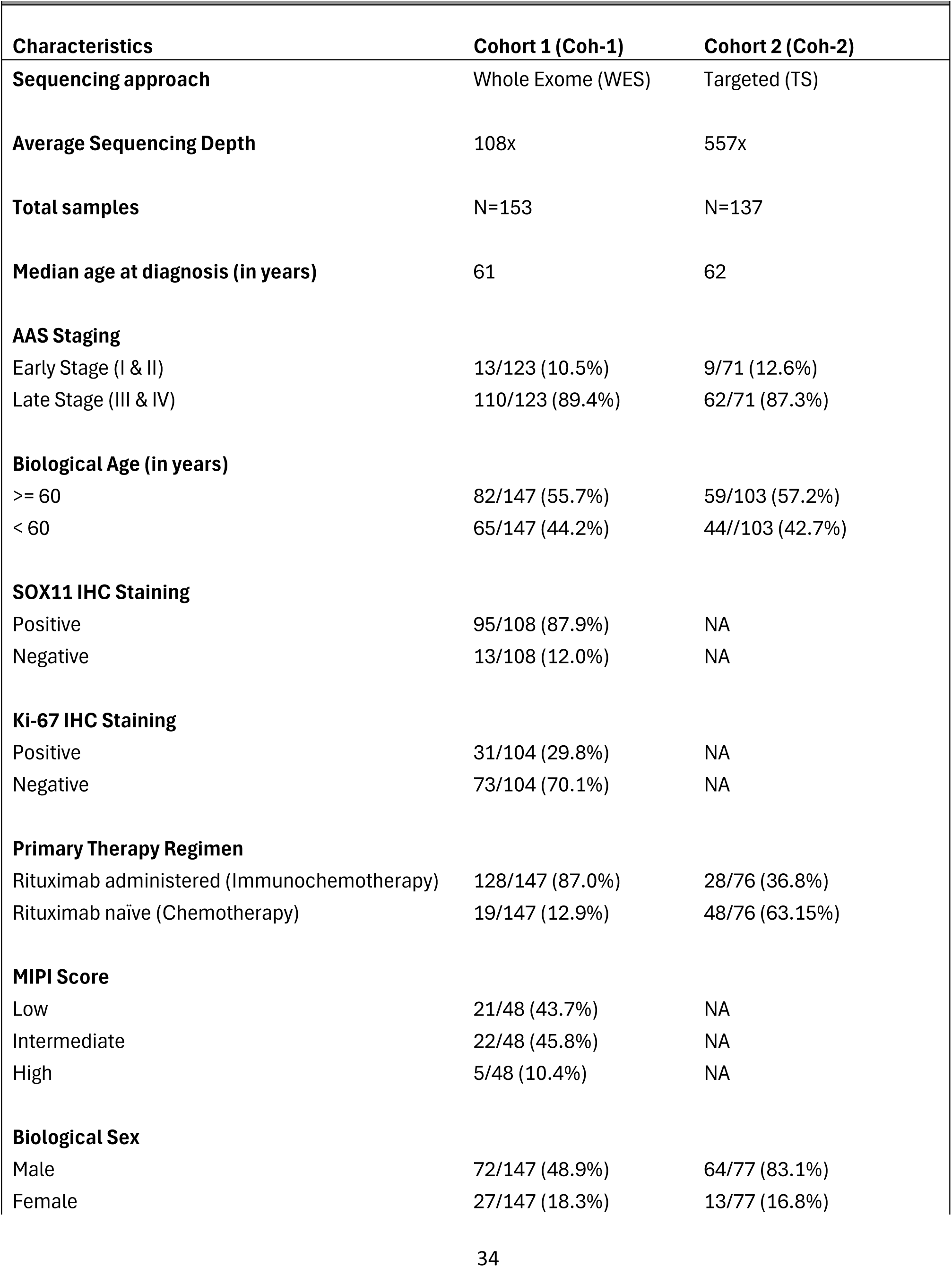
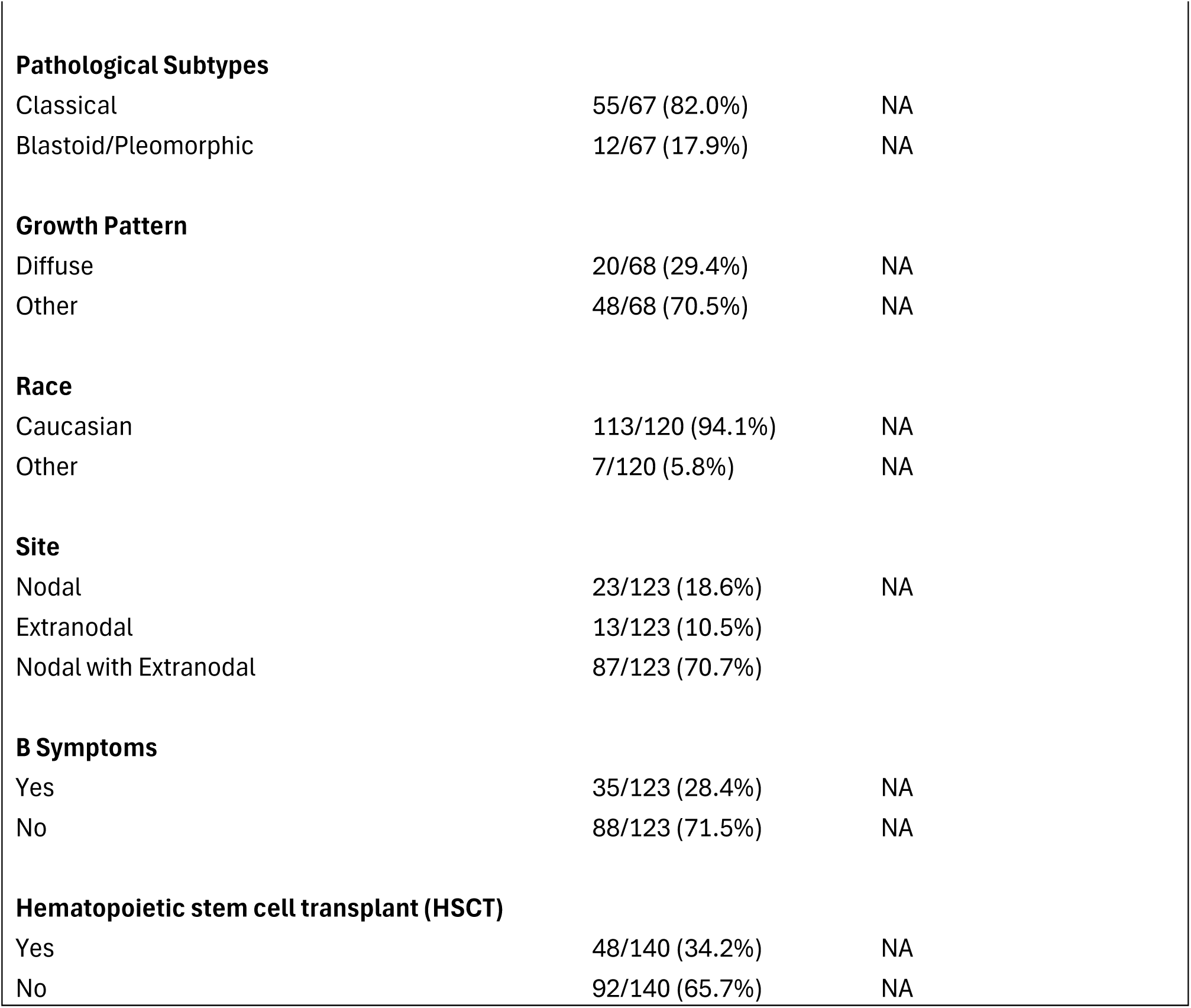
Clinical Characteristics of the Two MCL Cohorts.

Univariate analysis of Coh-1 showed that biological age, B-symptoms, and transplantation status was significantly associated with both overall survival (OS) and progression-free survival (PFS) (**Supplementary Fig. 1a**). In addition, P53 immunohistochemistry (IHC) positivity (score ≥ 90%) and absence of Rituximab in the primary treatment regimen were linked to poorer OS. Conversely, patients aged <60 years, without B symptoms, or who underwent transplantation exhibited improved OS and PFS (**Supplementary Fig. 1b, c**).

### Somatic mutation analysis alteration in genes regulating DNA repair and epigenome

WES analysis of Coh-1 identified 8,392 variants affecting 5,693 genes that passed the filtering criteria (**See Methods**). These primarily consisted of 8,374 single nucleotide variants (SNVs) and 558 insertions/deletions (**Supplementary Fig. 2a**). In Coh-2, we detected 1,775 SNVs and 110 insertions/deletions (**Supplementary Fig. 2a**). To validate SNV detection by WES, we also performed TS using a custom gene panel^17^ on 17 cases from Coh-1 and found an overlap of 94% of detected variants, which indicated 94% variants in TS could be recovered in WES with a correlation (r = 0.68, p < 0.05) of variant allele fraction (VAF) (**Supplementary Fig. 2b)**. The non-recoverable 6% of TS SNVs were mainly non-recurrent or rare alterations, likely influenced by high-depth coverage. As expected, mutational signatures indicative of known biological processes was observed, including SBS30 (defective base excision repair), SBS5 (age-related), and SBS4 (DNA damage from environmental mutagens), in Coh-1 (**Supplementary Fig. 2c**).

Coh-1 (n=153) revealed frequent mutations in DDR genes (*ATM*: 42%, *TP53*: 12%) and chromatin reorganization genes (*KMT2D*: 21%, *SMARCA4*: 13%, *SP140*: 10%, *ARID1A*: 9%, *KMT2C*: 8%, *KMT2B*: 5%) (**Fig. 1a**). Other frequent mutations included genes involved in cell cycle regulation (*CCND1*: 11%, *PCLO*: 5%, *SYNE1*: 7%), protein stability (*UBR5*: 11%), NF-κB signaling (*CARD11*: 8%, *NFKBIE*: 4%), apoptosis pathway (*BIRC3*: 5%, *SPTAN1*: 5%) and various other biological pathways (*LRP2*: 5%, *ANK2*: 5%, *SAMHD1*: 5%, *S1PR*1 *7*%, *FAT1*: 3%) (**Supplementary Table 2, 3**). Although most genes participate in multiple pathways, we have categorized them into a single pathway for simplicity (**Fig. 1a**). A similar mutational landscape was observed for Coh-2 (**Supplementary Fig. 3a, Supplementary Table 4**). Mostly, no significant differences in mutation frequencies of common genes were observed between the two cohorts, with a few exceptions, such as *NOTCH1*, *NOTCH2, LRP2, BCOR*, which showed higher detection rates in the TS (Coh-2) compared to WES (Coh-1). This increased sensitivity, attributable to greater coverage and depth in TS, suggests that these mutations are likely subclonal (**Supplementary Fig. 3b, Supplementary Table 5).**

**Fig. 1.**
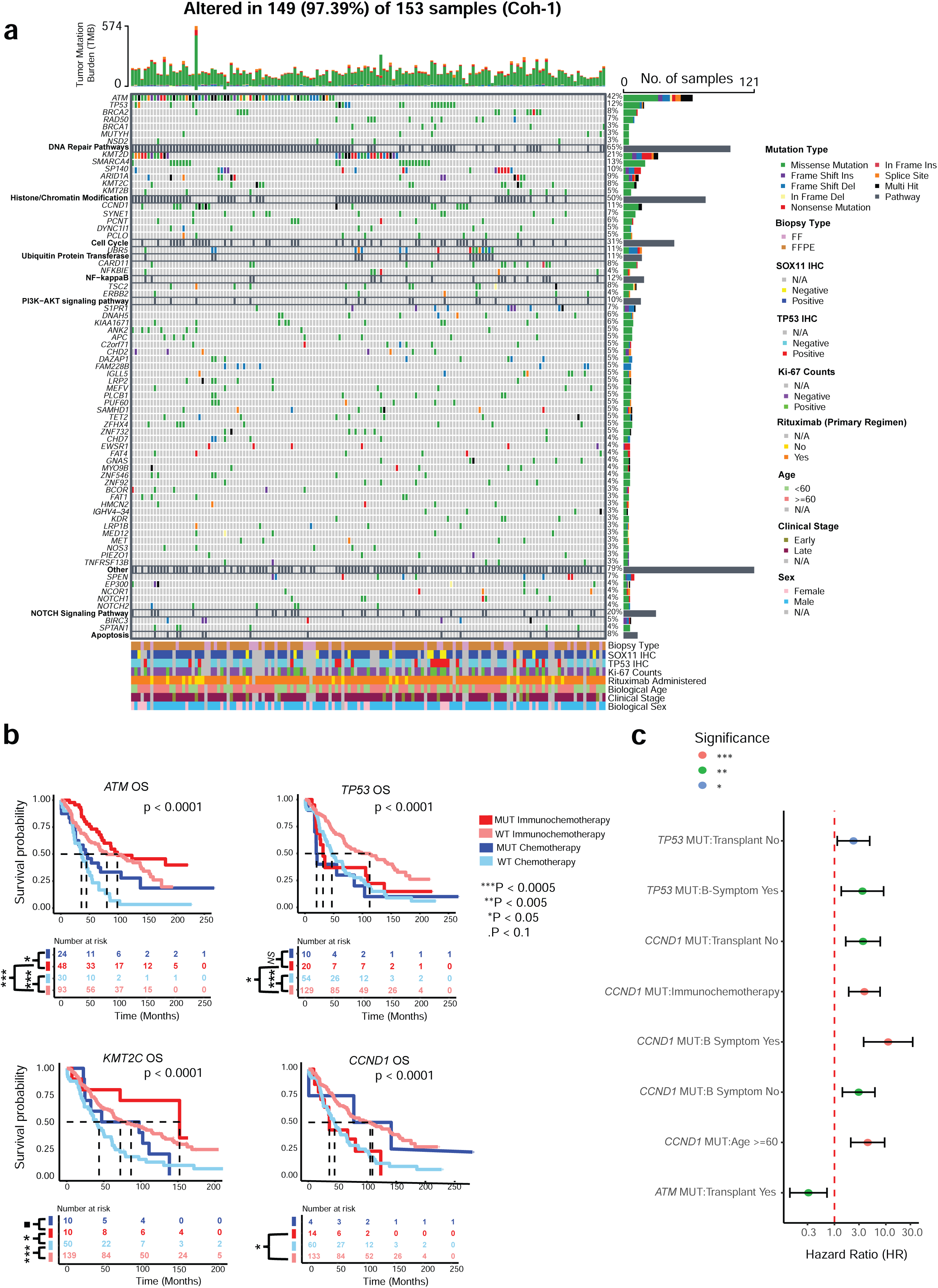
Genomic mutational landscape in MCL and associated clinical outcomes in Coh-1. **(a)** Oncoplot showing the frequency of recurrent genomic mutations (≥3%) in MCL patients (n = 153), with mutation types indicated by different colors. The right panel displays the percentage of each mutation type per gene. Clinical and pathological information for each patient includes primary treatment regimen (Immunochemotherapy: Rituximab administered vs. Chemotherapy: Rituximab naive), Ann Arbor stage, immunohistochemistry (IHC) status of TP53, SOX11, and Ki-67, source material type of the biopsy, biological age and sex. **(b)** Kaplan-Meier survival curves assessing OS, stratified by mutation status (presence/absence) of specific genes and primary treatment regimen (Immunochemotherapy vs. Chemotherapy). Patients from both Coh-1 and Coh-2 were combined for this analysis. The dotted black line represents median survival time (50% survival probability). Log-rank tests were used to compare survival curves across the four groups. ·P < 0.1, *P < 0.05, **P < 0.005, ***P < 0.0005. **(c)** Forest plot illustrating the hazard ratios (HRs) and 95% confidence intervals (CIs) for significant interaction terms between gene mutation status and clinical variables in relation to OS. Interaction terms include mutations in *ATM*, *TP53*, *KMT2C* and *CCND1* genes with clinical variable such as transplant status, presence of B symptoms, biological age, and primary treatment regimen (Immunochemotherapy vs. Chemotherapy). The vertical red dashed line indicates a hazard ratio of 1.0 (no effect). Colored points represent statistically significant interactions, with significance levels indicated by color: ***P < 0.0005 (red), **P < 0.005 (green), *P < 0.05 (blue).

There was a significant association between *TP53* mutations and positive p53 protein expression (IHC Score ≥ 0.9, r = 0.76, p < 0.05) (**Fig. 1a**). Hotspot mutation analysis in the combined cohort revealed mutations in the *TP53* are enriched in the DNA binding domain (DBD) whereas mutations in the *ATM* spanned its full length, with a novel hotspot missense mutation, R3008H, identified near the C-terminal PI3-kinase regulatory domain (**Supplementary Fig. 4a, b**). Hotspot mutations in *CCND1* (exon 1) such as V42A, Y44H/C, K46I/N, C47R, and K50E were identified (**Supplementary Fig. 4c**). While mutations in the 3’UTR of *CCND1* were less frequently detected in WES due to the region not being targeted in the capture, the TS revealed mutations in 4.7% of MCLs in the regions flanking the 3’UTR and 12% within the exonic regions (**Supplementary Table 6**).

Using combined OS data from both cohorts, we assessed how patients with and without gene mutations responded to chemotherapy and immunochemotherapy (**Fig. 1b**). For *TP53*, there was no survival difference between mutant (MUT) and wild-type (WT) cases following chemotherapy. However, the addition of immunochemotherapy improved outcomes in WT patients but not in those with T*P53* mutations, indicating that *TP53*-MUT MCLs does not benefit from immunochemotherapy. In contrast, for *ATM*, patients treated with immunochemotherapy, consistently showed better survival than those receiving chemotherapy alone, regardless of mutation status, highlighting that treatment modality had a greater impact on outcome than *ATM*-MUT status. Among *KMT2C* cases, the best outcomes were observed in mutant patients receiving immunochemotherapy. However, *KMT2C*-WT cases on immunochemotherapy showed similar survival to *KMT2C*-MUT cases on chemotherapy, suggesting mixed effects of mutation and treatment. Lastly, *CCND1*-MUT cases had worse survival even with immunochemotherapy. Although the chemotherapy group for *CCND1*-MUT was small, we concluded that *CCND1* mutations may be associated with poor prognosis despite immunochemotherapy.

To better understand how genetic mutations interact with clinical factors, we performed a multivariate Cox regression with interaction terms in Coh-1 (**Fig. 1c**). We focused on three clinical variables, such as biological age, B symptoms, and transplant status, all of which were individually associated with OS in univariate analysis. The analysis showed that *TP53***-**MUT patients had especially poor outcomes when they were not transplanted or had B symptoms. In contrast, *ATM***-**MUT patients appeared to benefit from transplant. *CCND1* mutations were associated with increased risk across all subgroups, with the strongest negative effect seen in patients with B symptoms. These results highlight how clinical and genomic features jointly shape survival even among patients receiving the same treatment, supporting more personalized risk stratification in MCL.

### Genomic DNA copy number analysis (CNA) profiling reveals key drivers of survival and treatment response

Comparative CNA analysis across two platforms (n=290) indicated similar aberrant genomic profiles with frequency differences in aberrant loci (**Fig. 2a, Supplementary Table 7**). Using GISTIC-2^18^ and iGC^19^, we identified focal gain (gain) lesions along with copy number driven genes respectively at 3q22.3 (37%: *DRB1, STAG1*), 3q23-3q29 (35-46%: *ATR, BCL6, DLG1*), 7p22.3 (16%: *RAC1, EIF3B*), 8q24.21 (20%: *MYC, ASAP1*), 15q21.2 (18%: *BCL2L10, FBN1, COPS2*), 18q21.3 (10%: *BCL2, TNFRSF11A*). Genes with downregulated expression linked to focal deletions (del) at 1p21.2 (42%: *AGL, DBT, CDC14A*), 6q21-6q25.3 (21%: *PRDM1, AIM1, ATG5*), 8p23.3 (27%: *ERICH1*), 9p21.1-21.3 (20-23%: *CDKN2A, CDKN2B*), 11q22.3 (38%: *ATM, CASP4, DDX10*), 13q12.11-13q14.2 (26%-40%: *LATS2, DLEU1*), 15q26.2 (15%: *MEF2A, RGMA*), 17p11.2-13.3 (17-30%: *TP53, DVL2, POLR2A*) and 19p13.3 (24%: *TLE2, MOB3A*) **(Fig. 2a, Supplementary Table 8,9)**. Integrative RNA-Seq and CNA analysis in Coh-1/Coh-2 revealed that upregulated genes were enriched in ATR, MYC, PI3K-AKT, and Wnt signaling pathways, whereas downregulated genes were associated with impaired DDR, NF-κB signaling, apoptosis, and cell cycle regulation **(Fig. 2b, Supplementary Table 10)**.

**Fig. 2.**
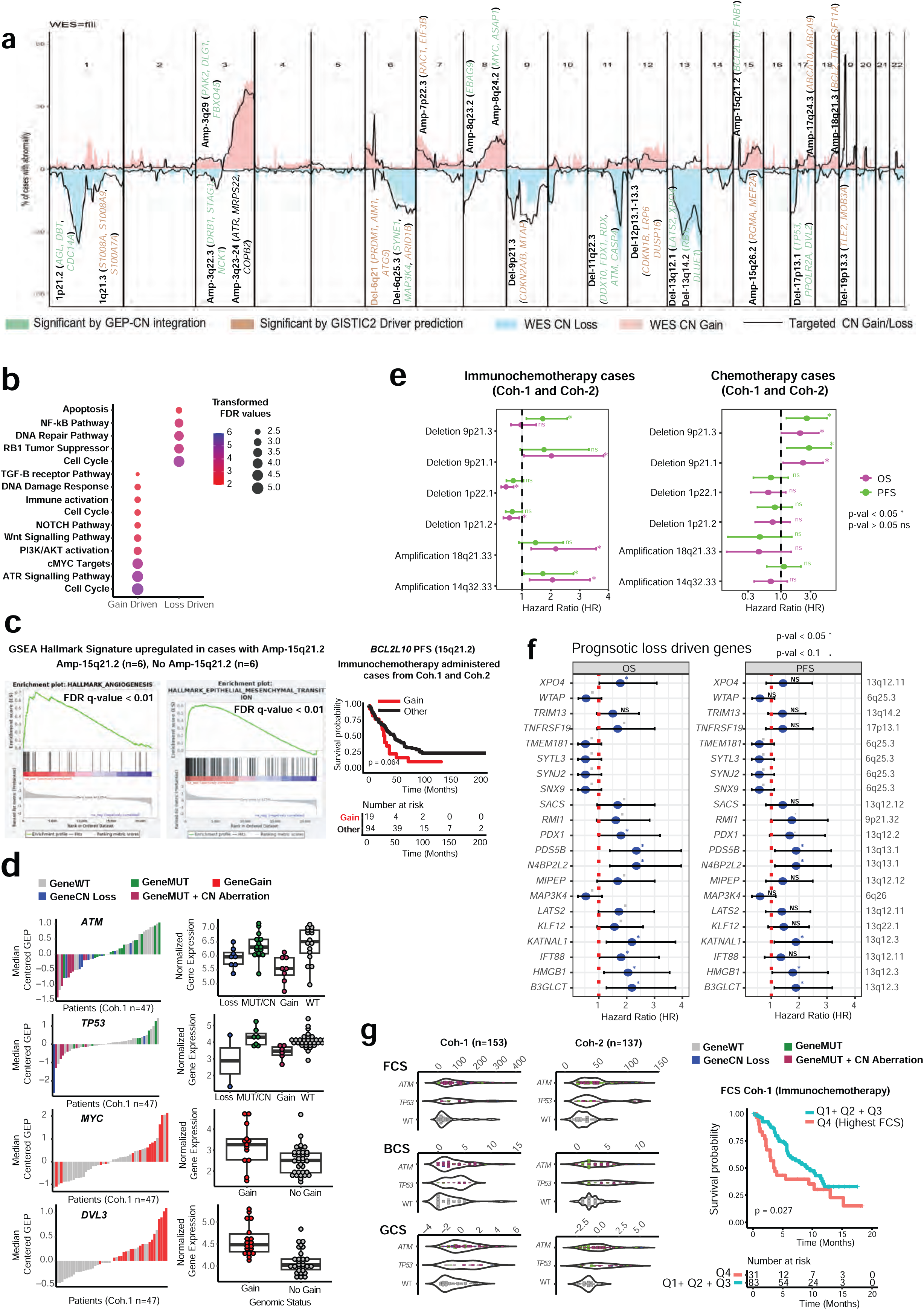
Comprehensive genomic copy number analysis (gCNA) in Coh-1 and Coh-2. **(a)** CNA frequency plot displaying genomic regions of gain (coral red) and loss (blue) in Coh-1, and corresponding changes in Coh-2 (solid black line). Gene annotations reflect both GEP and CNA status, with key drivers identified by GISTIC-2 across all 22 chromosomes. Lesions with adjusted p < 0.1 were included. **(b)** Pathway analysis of shared gain/loss-driven genes across both cohorts using ConsensusPathDB. Pathways with adjusted p < 0.05 were included. **(c)** GSEA comparing cases with and without Gain-15q21.2 (n=6 each) in Coh-1. *BCL2L10*, located within the amplified region, was identified as a potential driver and showed a trend toward poorer OS following immunochemotherapy. **(d)** Correlation between CNA status and mRNA expression of key driver genes (*ATM*, *TP53/MYC*, *DVL3*) in Coh-1. **(e)** Forest plots showing hazard ratios (HR) and 95% confidence intervals (CI) for OS (magenta) and progression-free survival (PFS, green) based on selected genomic alterations in patients treated with immunochemotherapy (left) or chemotherapy (right), using Cox models. Analyzed events include deletions (9p21.3, 9p21.1, 1p22.1, 1p21.2) and gains (18q21.33, 14q32.33). HR > 1 indicates increased risk; HR < 1 indicates reduced risk. *p < 0.05; ns = not significant. **(f)** Forest plots for HRs and 95% CIs of OS (left) and PFS (right) associated with loss of candidate driver genes, annotated with chromosomal loci. HR > 1 reflects increased risk; HR < 1 reflects potential protective effect. *p < 0.05; NS = not significant. **(g)** Violin plots showing distribution of copy number burden-focal (FCS), broad (BCS), and global (GCS) across *ATM*- and *TP53*-perturbed groups and wild-type (WT) cases in Coh-1 (n=153) and Coh-2 (n=137). Genomic alteration groups: WT (gray), mutation only (green), gain (red), loss (blue), and combined mutation + CNA (magenta). Kaplan-Meier curve (right) illustrates survival based on FCS burden, comparing Quartile 4 (Q4): Highest FCS: salmon vs. combined Q3+Q2+Q1: cyan groups.

We observed a novel gain at 15q21.2 in 18% of cases (52/290), with *BCL2L10* emerging as a potential driver (**Fig. 2c**). Gene Set Enrichment Analysis (GSEA) of RNA-Seq data from Coh-1 comparing cases with gain-15q21.2 (n=6) to WT-15q21.2 (n=6) revealed significant enrichment of gene signatures related to angiogenesis and epithelial-mesenchymal transition (EMT) (FDR q < 0.05, **Supplementary Table 11**). Gain-15q21.2 was associated with a trend toward inferior PFS in immunochemotherapy-treated patients, though no difference in OS was observed (**Fig. 2c**). As expected, the mRNA expression levels of commonly known drivers such as *ATM*, *TP53*, *MYC*, and *DVL3* were significantly correlated with their corresponding genetic aberrations (**Fig. 2d**). Survival analysis showed that del-9p21.3 was linked to poor prognosis regardless of treatment, though immunochemotherapy demonstrated improved outcomes versus chemotherapy alone. In contrast, gain-14q.32.33 showed no significant benefit from immunochemotherapy (**Fig. 2e**). Multivariate Cox regression analysis using interaction terms in Coh-1 showed gain-13q31.3 were associated with increased risk in non-transplanted and older patients, while del-6q21/del-1q21.1 showed a protective effect in patients treated with immunochemotherapy. Multiple gains on chromosome 8q were associated with improved outcomes in the context of Rituximab treatment or B symptoms highlighting the combined influence of genomic and clinical factors on MCL survival (**Supplementary Fig. 5**).

We identified a prognostic loss-driven gene signature from 6q (*WTAP*, *MAP3K4*, *SYTL3*, *SNX9*) and 13q (*LATS2*, *HMGB1*, *IFT88*, *PDX1*) in immunochemotherapy-treated Coh-1 cases. (**Fig. 2f**). MCLs with dual loss (i.e. mutation and deletion of *ATM* or *TP53*) demonstrated significantly higher broad, focal, and global CNA scores (BCS, FCS, and GCS, respectively) than those with single alterations or WT cases. Notably, higher CNA burden is associated with a poor prognosis in immunochemotherapy-treated cases (**Fig. 2g, Supplementary Table 12).** These findings highlight the prognostic relevance of CNA-driven alterations and underscore the impact of CNA on treatment outcomes in MCL.

### Cohort-guided Non-Negative Matrix Factorization (NMF) consensus clustering reveals robust, clinically relevant prognostic subgroups

Somatic interaction analysis was performed across Coh-1 and 2 (n=290), using genomic events with comparable frequencies to identify co-occurring alterations (**Supplementary Fig. 6**). *TP53* mutations co-occurred with del-13q14.2, del-17p13.3, del-9p21.1-21.3, del-6q21, and *ARID1A* mutations, while *ATM* mutations co-occurred with del-13q14.2, del-11q22.3, and del-6q25.3. *CCND1* mutations co-occurred with del-13q14.2 and gain-15q21.2. Given the broader genomic coverage, clinical annotation, and immunochemotherapy predominance of Coh-1, we applied NMF consensus clustering on this cohort using 35 recurrent alterations observed in more than 3% of cases (**Supplementary Table 13)**, which identified six distinct molecular clusters (C1-C6) (**Fig. 3a**, **Supplementary Table 14**). Key genetic characteristics of C1-C6 clusters are detailed in **Table 2**. Kaplan-Meier analysis showed distinct 3-year OS patterns: C5 had the worst survival (38.5%), while C1 and C6 showed the best (93.3% and 94.4%) (**Fig. 3b, left**). These clusters differed by clinical features, including p53 IHC enrichment in C5, female predominance, and low blastoid/pleomorphic morphology in C1 (**Supplementary Table 15**). For simplicity, the six clusters were grouped into high-risk (C5), intermediate-risk (C2-C4), and low-risk (C1, C6) subgroups, each with distinct OS profiles (**Fig. 3b, right**). GSEA analysis of transcriptomic data revealed significant enrichment of the proliferation-promoting gene signatures, including MYC, G2M, E2F, and MTORC1 enrichment in the “high-risk”, EMT/angiogenesis in the “intermediate-risk”, and immune activation in the “low-risk” clusters (**Fig. 3c**). FCS differed significantly between low- and intermediate-risk groups, while BCS and GCS showed no significant differences across clusters (**Supplementary Fig. 8a**). CIBERSORTx analysis showed elevated memory B cells in the “high-risk” MCL, while T_FH_ and CD8^+^ T cells were significantly (p < 0.1) enriched in “low-risk” cases (**Fig. 3d, Supplementary Fig. 8b**).

**Fig. 3.**
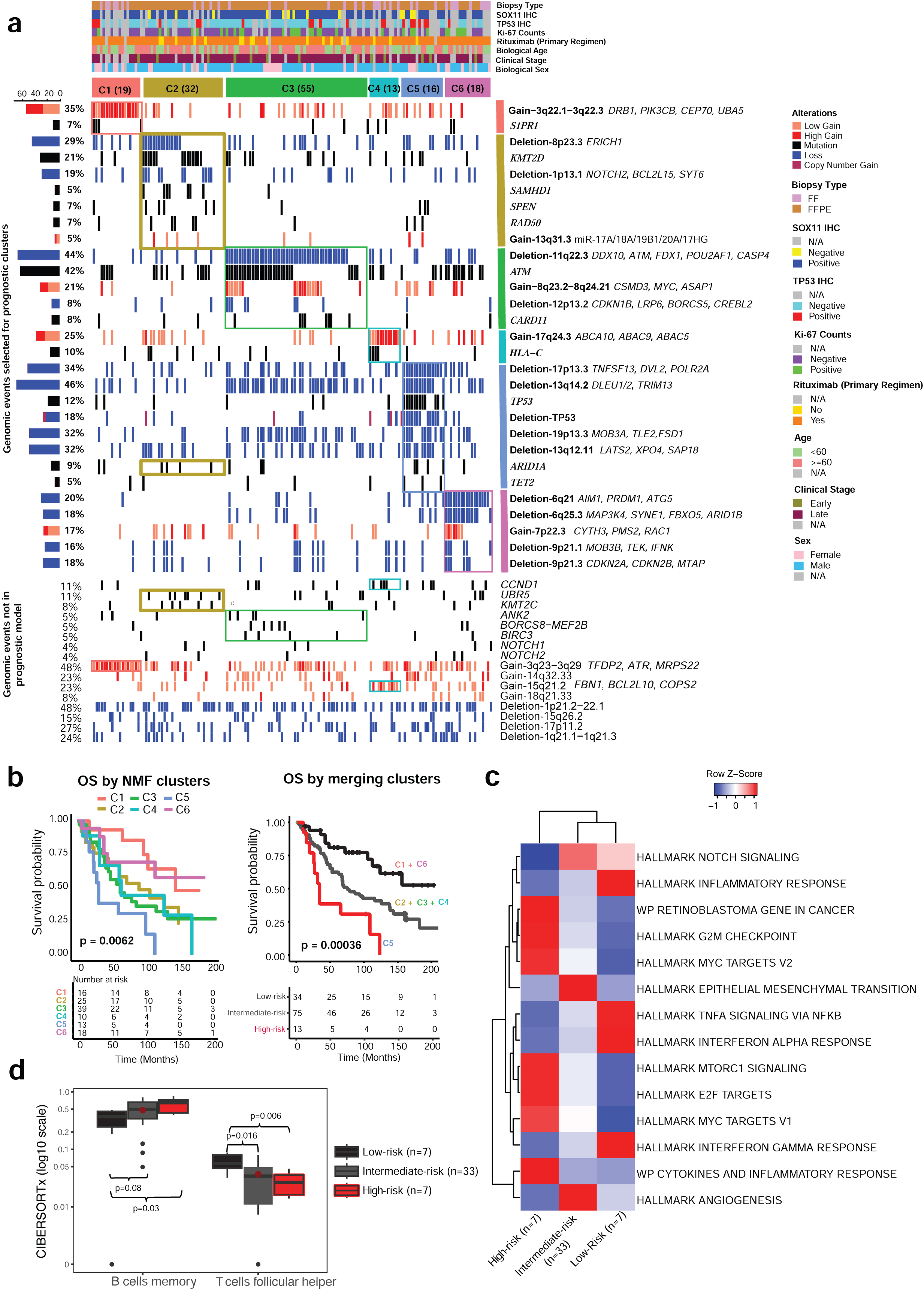
Genomic landscape of the six prognostic clusters identified in Coh-1. **(a)** Heatmap shows the distribution of recurrent somatic alterations across six prognostic clusters (C1– C6) derived using Non-Negative Matrix Factorization (NMF) consensus clustering. Top annotations represent clinical and pathological features, including biopsy type, SOX11 and TP53 IHC status, Ki-67 index, rituximab treatment, age, clinical stage, and sex. Rows correspond to genomic events: copy number deletions (blue), gains (red), and gene mutations (black) with events either selected (top panel) or not selected (bottom panel) for the prognostic model. Cluster-specific enrichment patterns highlight distinct molecular subtypes, with bolded genes indicating key alterations contributing to prognosis. Left-side bar plots show alteration frequencies across the cohort. **(b)** Kaplan-Meier survival curves depict overall survival (OS) stratified by NMF consensus clusters (left) and merged prognostic groups (right). In the left panel, OS is shown for all six NMF clusters (C1-C6), revealing significant survival differences across subgroups (*p* = 0.0062). In the right panel, clusters were merged based on OS patterns into three prognostic categories: low-risk (C1 + C6, red), intermediate-risk (C2 + C3 + C4, green), and high-risk (C5, blue), showing enhanced separation of survival outcomes (*p* = 0.00036). The number at risk over time is indicated below in the strata table. These merged groups highlight the prognostic utility of molecular clustering for survival stratification. **(c)** Gene Set Enrichment Analysis (GSEA) on RNA-seq expression profiles in Coh-1 to compare pathway-level activity across cases stratified by Low-risk, Intermediate-risk and High-risk groups. The heatmap displays the average row Z-scores of MSigDB Hallmark and WikiPathways gene signatures for each survival group, capturing relative pathway activation. **(d)** CIBERSORTx-based bulk immune profiling of T-cells (follicular helper) and B Cell Memory across overall survival-defined prognostic groups (high-risk, intermediate-risk, low-risk). Statistical comparisons were performed using the Wilcoxon rank sum test; significance levels were annotated as “ns” for non-significant comparisons.

**Table 2.** Genetic Subtype Overview of MCL.

| Cluster | Subtype Designation | Characteristics |
| --- | --- | --- |
| C1 | Amp3q | Amp3q (*BCL6*, *DBR1*, *PIK3CB*, *ATR*) |
| C2 | Epigenetic Modifier-Enriched | Del 8p23.3<br>KMT2D/KMT2C mutations<br>SPEN mutations<br>SAMHD1 mutations<br>RAD50 mutations<br>UBR5 mutations |
| C3 | *ATM* inactive | Del11q22.3 (*CASP4*, *ATM*, *DDX10*)<br>*ATM* mutations<br>Amp 8q23-24 (*MYC*, *ASAP1*) |
| C4 | Amp17q | Amp17q24.3 (*ABCA9*)<br>Amp15q21.2 (*FGF7*, *BCL2L10*) |
| C5 | *TP53* inactive | Del17p<br>*TP53* mutation<br>Del13q14.2 (*RB1*, *DLEU1*, *DLEU2*)<br>ARID1A/TET2 mutations |
| C6 | Del6q | Del6q (*SYNE1*, *PRDM1*, *MAP3K4*) |

To extend subtype analysis across cohorts, we applied a pan-cohort framework using a random forest classifier trained on Coh-1 to assign Coh-2 samples to existing clusters, leveraging its robustness to sparse genetic events resulting from differences between WES (Coh-1) and TS (Coh-2) (**Supplementary Table 16**). Five of six clusters were recovered, with no cases assigned to C4, likely due to missing *HLA-C* mutation data and underrepresentation of gain-17q24.3 on TS (Coh-2). Across the full dataset, the “high-risk” group (10.6%) was marked by *TP53* mutations and co-deletions in 17p13.3, 13q14.2, 19p13.3 (**Fig. 4a**). The “intermediate risk” group (65.7%) was enriched for *ATM* mutations, del11q, 8q/15q/17q gains, and epigenetic regulator mutations (*KMT2D*, *ARID1A*, *UBR5*). The “low-risk” group (24.1%) lacked biallelic *TP53* inactivation and showed isolated *ATM* mutations without 11q loss, suggesting monoallelic or less disruptive events. Survival analysis confirmed stratification by these clusters, with C5 showing the worst, and C1/C6 the most favorable OS (**Fig. 4b, left**). Merging into three risk groups improved prognostic separation (log-rank p < 0.0001, **Fig. 4b, right),** highlighting the biological and clinical relevance of this classification (**Fig. 4c**).

**Fig. 4.**
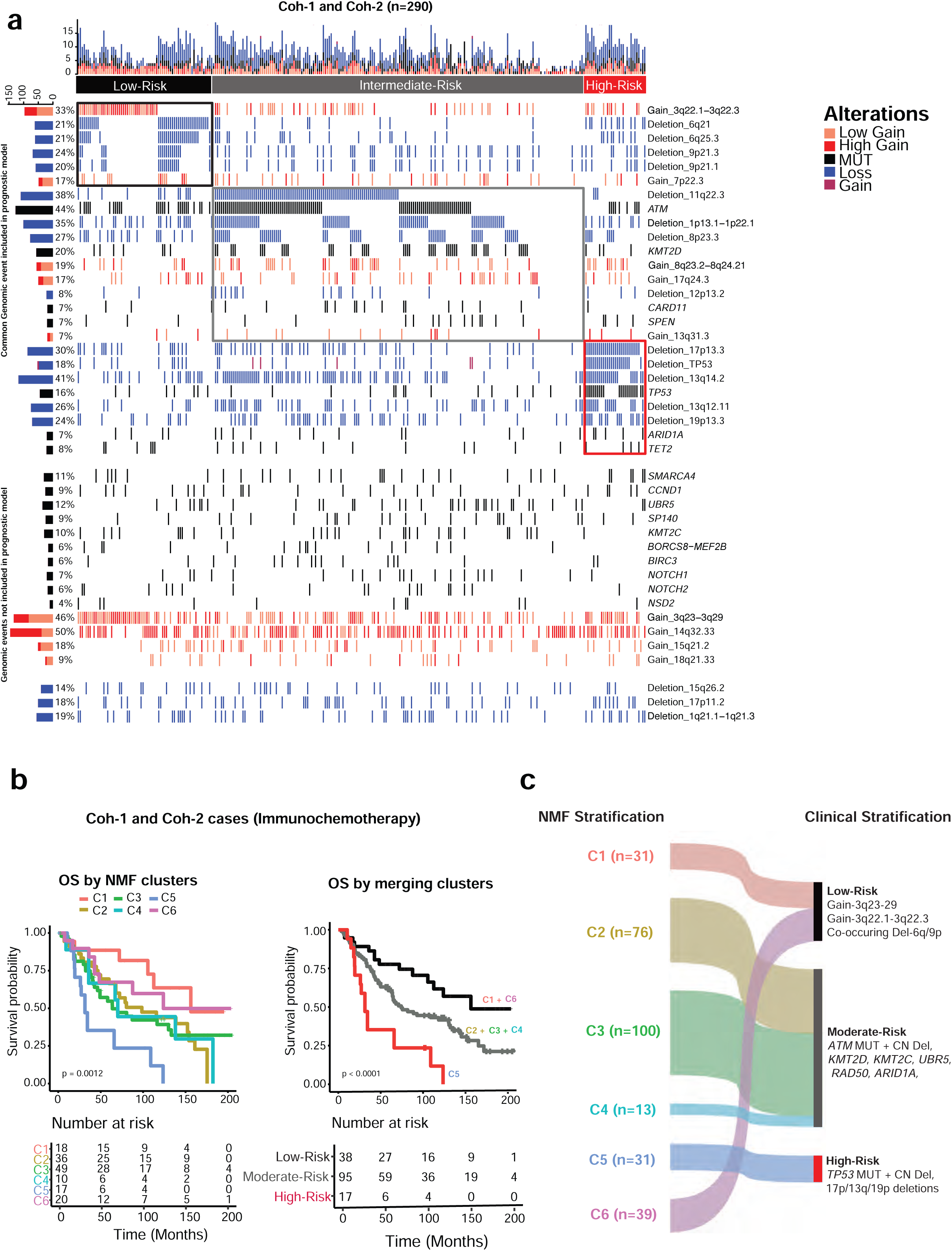
Validation of NMF-derived genetic clusters across Coh-1 and Coh-2. **(a)** Oncoplot of genomic alterations in 290 patients (columns), annotated by merged prognostic groups: low-risk (C1+C6), intermediate-risk (C2+C3+C4), and high-risk (C5). Rows represent genomic events (mutations, gains, deletions), with those used in the original NMF model (from Coh-1) shown in the upper section. A visual gap separates these from additional genetic events not included in the clustering algorithm but retained to provide broader genomic context. **(b)** Kaplan-Meier curves comparing OS by NMF-defined clusters (left) and merged prognostic groups (right), in patients receiving immunochemotherapy. Log-rank test was used for significance. **(c)** Sankey diagram showing the correspondence between NMF clusters (C1-C6) and three clinically stratified prognostic groups. Stream width reflects the number of cases transitioning from molecular clusters to broader clinical categories based on shared genomic features and survival patterns.

### Contrasting immune and spatial landscapes distinguish *TP53*- and *ATM*-perturbed MCLs

To delineate the cellular immune phenotypes and spatial TME heterogeneity linked to genetic subtypes, IMC analysis was conducted using a 36-biomarker panel **(Supplementary Fig. 9a, Supplementary Table 17)**. For IMC phenotyping analysis, a total of 50 samples were included: 42 MCL cases with sequencing data from Coh-1 and Coh-2 classified into *ATM*-perturbed (n=22), *TP53*-perturbed (n=14), and (*ATM*/*TP53*)-WT (n=6) along with 5 MCL cases without sequencing and 3 normal tissues (**Supplementary Fig. 9b**, **Supplementary Table 18**). Using PhenoGraph^20^ and a customized analysis (**Supplementary Fig. 9c**), we identified nine major cellular phenotypes that capture the complexity of the MCL TME (**Fig. 5a**). These phenotypes were functionally annotated based on the expression of activation, immunosuppressive, and tumor-associated markers (**Fig. 5a, Supplementary Table 19**). IMC images from three representative cases are presented, displaying phenotype maps alongside high-resolution marker expression channels at 50 µm scale (**Fig. 5b**). Among all MCLs analyzed, the most abundant populations were CD20⁺CCND1⁺SOX11⁺ (32.9%) and CD20⁺CCND1⁺SOX11⁻ (27.4%) tumor cells (**Fig. 5c**). Within the TME, CD3⁺CD8⁺ T cells represented the largest population (11.7%), followed by myeloid cells (11.3%), which included both intermediate (CD68⁻, 9.0%) and fully differentiated macrophages (CD68⁺, 2.3%). Importantly, both myeloid subsets expressed CD163, with particularly high levels in CD68⁺ macrophages, suggesting a polarization toward an M2-like (pro-tumor) phenotype^21,22^. CD3⁺CD4⁺ T cells accounted for 9.6%, while FOXP3⁺ regulatory T cells (Tregs) represented 4.0% of the TME. The least abundant populations included CD31⁺ endothelial cells (2.5%) and CD11c⁺ dendritic cells (0.7%). Notably, immune checkpoint markers including PD-1, TIM-3, and LAG-3 were highly expressed on CD8⁺ T cells, indicating functional exhaustion and impaired anti-tumor immunity (**Fig. 5a**). To ensure uniformity across tissue sources (two TMAs and unstained slides), we applied cyCombine for batch correction. Patients from different tissue sources and cohorts were well interspersed, with no batch effects and no differences in IMC cell fractions by tissue source origin respectively (**Supplementary Fig. 9b, 9d**).

**Fig. 5.**
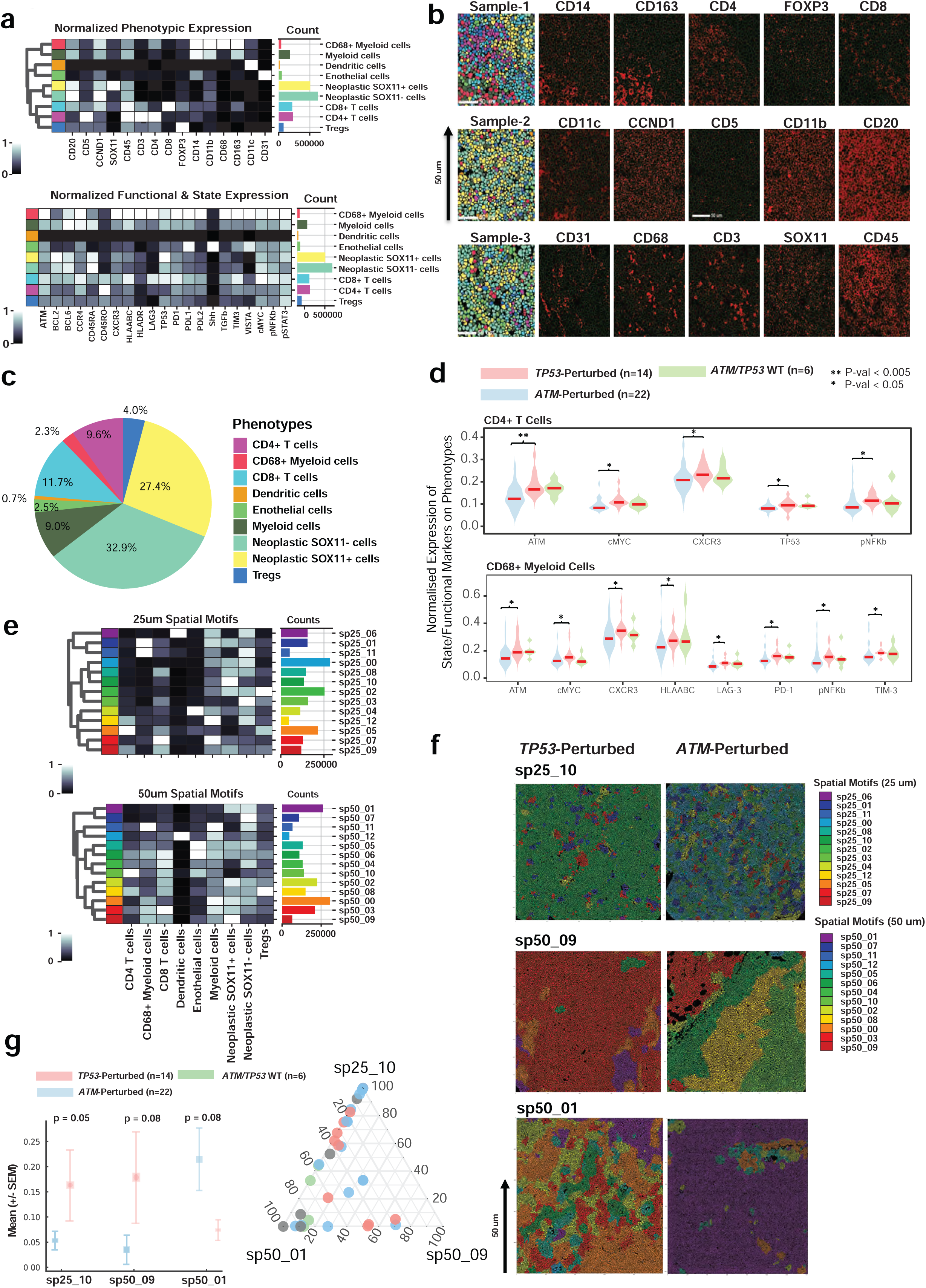
TME analysis using multi-proteomics IMC profiling. **(a)** Heatmaps of normalized expression (Z-scores, 0-1 scale) for canonical immune/tumor (top) and functional/state (bottom) protein markers across single cells from ROIs in all patients. Cell phenotypes were defined based on average marker expression. Bar graphs indicate cell counts per phenotype. **(b)** Representative multiplex images (three samples) show segmented ROIs with selected immune, tumor, and microenvironment markers. Markers are shown in red, nuclei in grayscale. Scale bars = 50 μm. **(c)** Pie chart showing the average proportions of major immune and tumor phenotypes across all samples, classified by canonical immune and tumor markers expression. **(d)** Violin plots showing differential expression of functional and state markers in CD4⁺ T cells (top) and CD68⁺ myeloid cells (bottom) across genetic perturbation groups. Medians shown as red bars; Statistical comparisons were performed using the Kruskal-Wallis test. **(e)** Spatial motif heatmaps illustrating co-localization patterns among immune and tumor phenotypes at 25 μm (top) and 50 μm (bottom) radii. Columns denote cell types; rows denote spatial motifs. Color intensity reflects normalized frequency (0-1). Bar plots (right) show absolute motif counts across samples. **(f)** Spatial maps of representative ROIs from *TP53*- and *ATM*-perturbed samples, highlighting differential motif composition at 25 μm and 50 μm radii. Color-coded motifs reflect distinct spatial arrangements of immune and tumor cells. **(g)** Left: Mean proportions (± SEM) of three spatial neighborhoods states (sp25_10, sp50_09, sp50_01) across genetically perturbed groups. Right: Ternary plot visualizing the relative contributions of the three spatial neighborhoods across all MCLs.

To better characterize immune dynamics in MCL, we compared immune phenotypes across *TP53*-perturbed, *ATM*-perturbed, and WT cases. Notably, *TP53*-perturbed MCLs displayed elevated expression of CXCR3, pNF-κB, and cMYC in CD4⁺ T cells (**Fig. 5d**). The overall myeloid cell population was increased in *TP53*-perturbed cases (**Supplementary Fig. 8e**), and CD68⁺ myeloid cells exhibited higher levels of immune checkpoint molecules, including LAG-3, TIM-3, and PD-1, indicating an immunosuppressive phenotype. Concurrent upregulation of cMYC and pNF-κB in these cells further suggests a state of heightened inflammatory signaling and metabolic activity. Collectively, these findings support the presence of an immune-infiltrated (’immune-hot’) yet functionally suppressed TME in *TP53*-perturbed MCL. Furthermore, we performed spatial neighborhood analysis at 25 µm and 50 µm across 50 MCL samples and identified 13 unique spatial neighborhoods each (**Fig. 5e**). Three of these significantly distinguished *TP53*- from *ATM*-perturbed tumors (**Fig. 5f, g**). sp25_10, enriched in SOX11⁺ tumor cells, and sp50_09, characterized by interactions among SOX11⁺ tumor cells, CD68⁺/⁻ myeloid cells, and CD8⁺ T cells, were significantly increased in *TP53*-perturbed MCLs (p < 0.1). In contrast, sp50_01, a mixed SOX11⁺/⁻ tumor region, was more frequent in *ATM*-perturbed cases. These findings suggest a more homogeneous TME in *TP53*-perturbed MCL driven by expansion of a dominant SOX11⁺ clone, while *ATM*-perturbed tumors maintain spatial heterogeneity. Additionally, *TP53* perturbation appears to influence myeloid cell localization around tumor clusters, potentially shaping an immune-modulated microenvironment. Collectively, these results underscore the complex spatial and functional architecture of the MCL TME.

### *TP53* loss leads to enhanced BCR signaling

To delineate the role of p53, we utilized four MCL cell lines for *TP53* modification **(Fig. 6a)**. Initially, we introduced WT-*TP53* into two *TP53*-deficient MCL cell lines, Maver-1 and Mino, in a doxycycline (Dox)-inducible approach and performed single-clone selection for the modified cells. The expression of WT-*TP53* was detrimental to the cells as minimal induction caused a significant cytostatic effect, whereas more intense induction induced substantial apoptosis (**Supplementary Fig. 10a**). Next, we generated *TP53* deficiency in the Z-138 cell line (WT-*TP53*) by creating either knockout (KO) or two hotspot mutations, specifically *TP53^R248Q^* and *TP53^R273C^* (**Supplementary Table 20**). Given the inherently challenging nature of transfecting lymphoma cells, we developed all-in-one vectors, containing high-fidelity SpCas9^23^ or Cas9-deaminase^24,25^ to facilitate the generation of the *TP53^KO^* or mutant cell lines. Through single clone selection, we successfully obtained pure clones with either KO or mutation (**Supplementary Fig. 10b-d**). For the p53 functional profiling, we first performed RNA-Seq in all the isogenic pairs. As expected, canonical p53 targeted genes, such as *CDKN1A, BBC3, MDM2,* and *BAX*, were shown to be substantially upregulated upon WT-*TP53* activation **(Fig. 6b, Supplementary Table 21)**. GSEA analysis revealed that activation of WT-*TP53* affects not only the cell cycle, DDR, and *TP53* signaling pathways but also significantly represses BCR signaling **(Fig. 6c)**. CUT&RUN analysis revealed that mutant-*TP53* exhibits a greater DNA binding abundance than WT-*TP53*, especially in Z-138 cells that express only WT-*TP53*. This suggests that mutant-*TP53* possesses a broader binding spectrum, potentially at the expense of its normal tumor suppressor functions **(Fig. 6d)**. Gene promoters differentially bound between WT and mutant-*TP53* are enriched in known *TP53* targets and novel ones, including *CCNE1*, *PCNA*, *SYK,* and *BCL2* **(Fig. 6e, f).** Upregulated pathways, encompassing p53 signaling, DDR, and PI3K-Akt signaling, were significantly enriched in WT-*TP53* compared to mutant-*TP53* cases, highlighting enhanced activation of tumor suppressive and immune-related programs **(Fig. 6g)**.

**Fig. 6.**
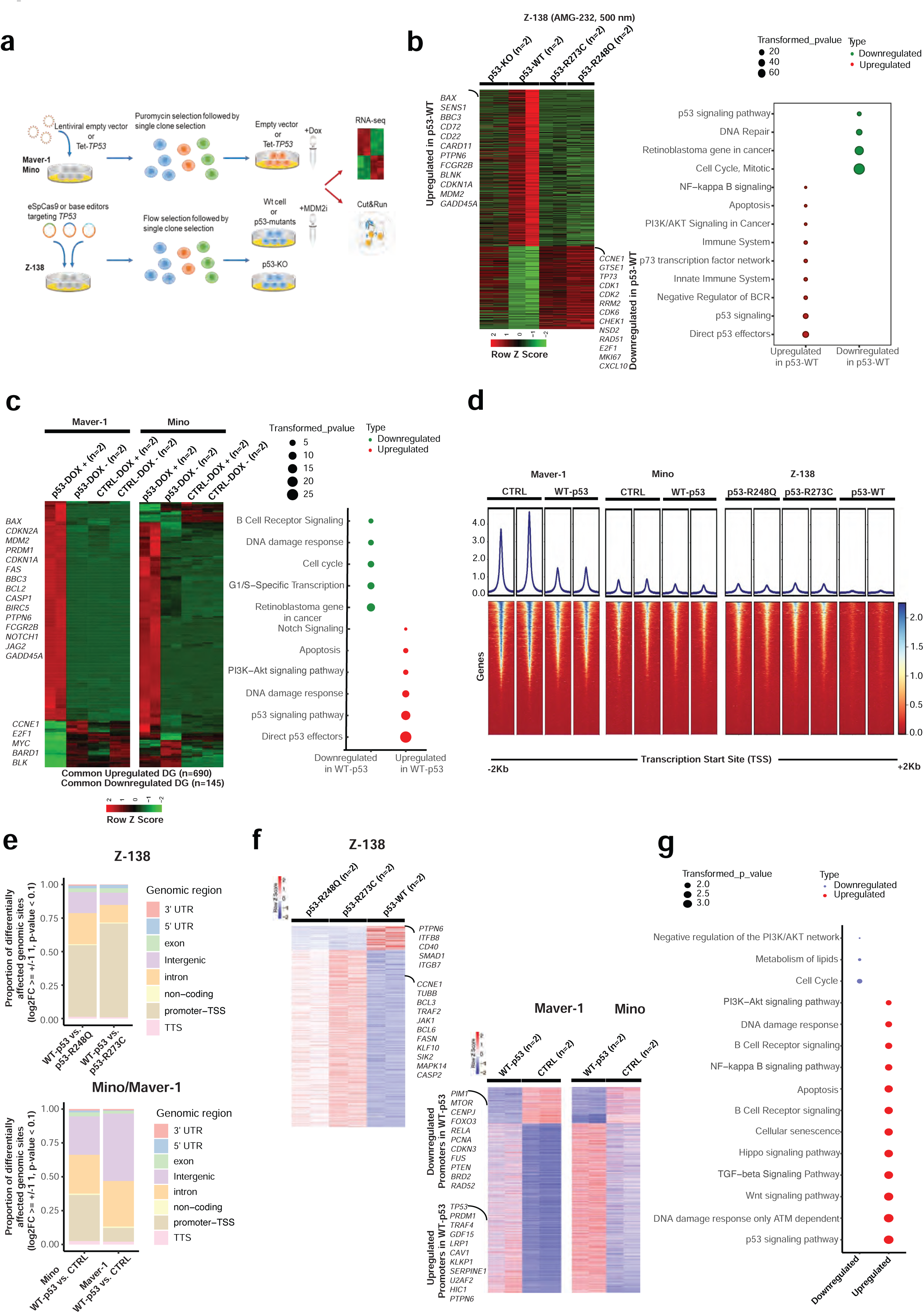
Regulation of signaling pathways by *TP53* in MCL. **(a)** Diagram detailing the development of *in vitro* models used to investigate TP53 function in MCL. **(b)** RNA-Seq analysis identifying DEGs and their functions following TP53 activation. Z-138 isogenic cell pairs were treated with the MDM2 inhibitor AMG-232 (500 nM, 24 h) to pharmacologically activate TP53. **(c)** In Maver-1 and Mino isogenic cell models, *TP53* was induced using doxycycline (125 nM for Maver-1; 250 nM for Mino, 24 h). **(d)** Overview of CUT&RUN assay showing TP53 binding distribution relative to transcription start sites (TSS). **(e)** Proportional distribution of differentially affected genomic regions across mutant p53 cell lines. Bar plots represent the proportion of genomic sites with significant changes (log₂FC ≥ ±1, p-value < 0.1) across annotated genomic features in Z-138 (p53-R248Q, p53-R273C) and Mino/Maver-1 cell lines. **(f)** Heatmap displaying gene targets affected by differential binding of WT-versus mutant TP53 as analyzed by CUT&RUN analysis. Each assay was performed using duplicates. **(g)** GSEA analysis illustrating pathway enrichment of genes directly influenced by p53 binding in Maver-1 and Mino cells with enforced WT-*TP53* expression.

### p53 represses BCR signaling through upregulating PTPN6 expression

We further explored BCR signaling regulation by *TP53* from the above findings, given the poor clinical response of *TP53* aberrant MCL to R-CHOP or BTK inhibitor treatment^26–28^. Since several BCR proximal signaling components, including both activators and antagonists, were regulated by p53, to determine the overall impact of *TP53* on BCR signaling, we induced the WT-*TP53* expression in *TP53*-deficient MCL cell lines and assessed the resulting BCR signaling activity. Notably, we observed a marked reduction in BCR signaling activity upon ectopic p53 expression, as indicated by decreased phosphorylation of SYK, BTK, and BLNK (**Fig. 7a**). To exclude the confounding effects of caspase activation, we treated the cells with the pan-caspase inhibitor Z-VAD-FMK prior to WT-*TP53* induction, which also resulted in reduced BCR signaling (**Supplementary Fig. 12a**). Consistently, when *TP53* is activated in the Z-138 isogenic pairs, BCR signaling notably reduced in the WT cells, whereas it remained unchanged in KO and homozygous mutant cells (**Supplementary Fig. 12b**). Interestingly, in the heterozygous mutant cells, we also noted a decrease in BCR signaling activity, albeit to a lesser degree than in WT cells, which further underscores the close association between BCR signaling and functional p53 levels **(Supplementary Fig. 12b)**.

**Fig. 7.**
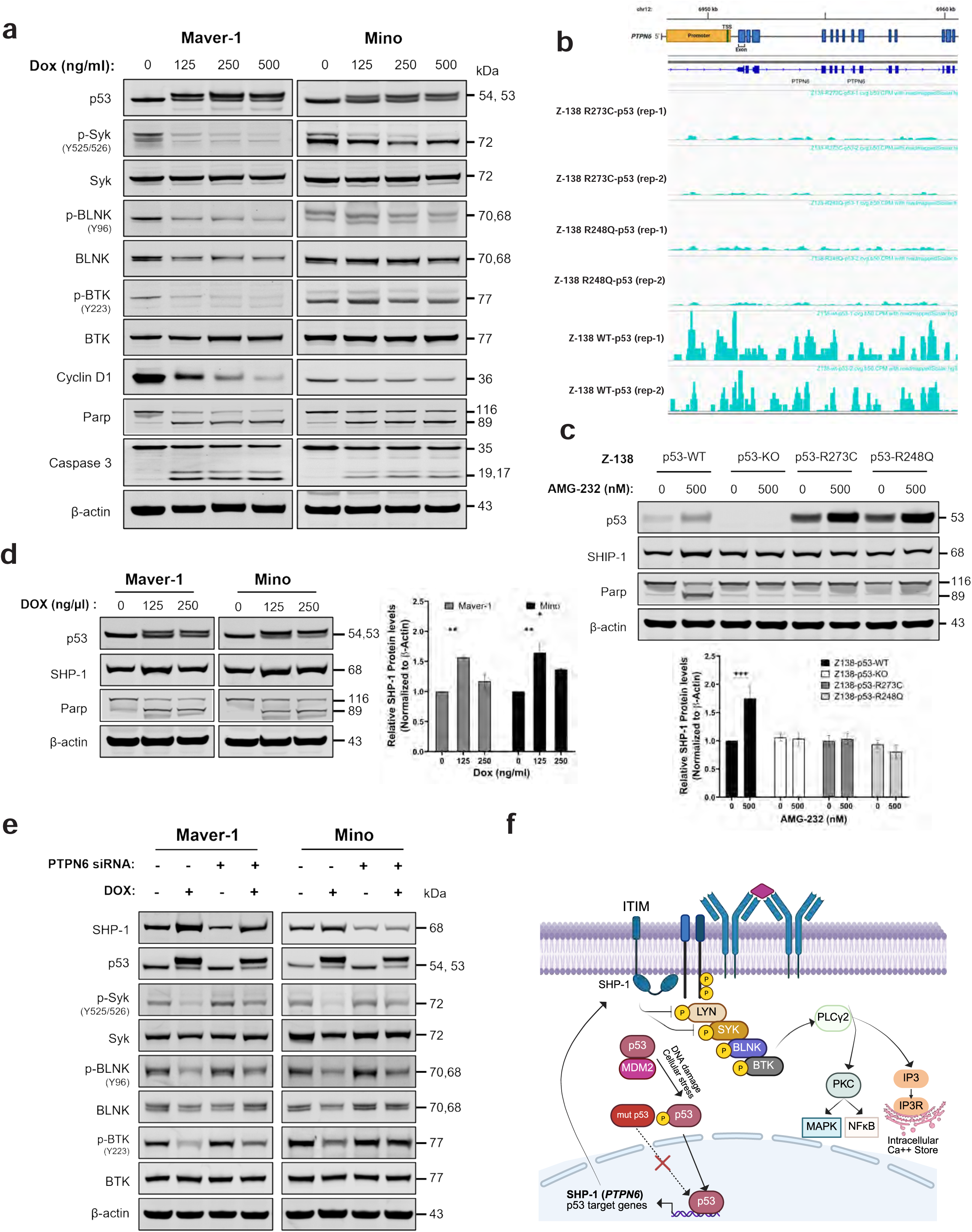
P53 suppression of BCR signaling via upregulation of PTPN6. **(a)** Examination of BCR signaling and Cyclin D1 level by WB in Marver-1 and Mino cells upon enforced TP53 expression. **(b)** CUT&Run data results demonstrating the binding differences between wt- and mutant-p53 within the *PTPN6* gen area. **(c and d)** Validation of PTPN6 (SHP-1) upregulation by p53 in Z-138 (C) and Maver-1/Mino isogenic pairs upon p53 activation by WB. Adjacent column charts quantify SHP-1 levels by normalizing SHP-1 signal pixel-intensity to β-actin. **(e)** Examination of BCR signaling pathway following PTPN6 knockdown (KD). The modified Maver-1 and Mino cells were KD with siRNA for 48h followed by treatment of Dox treatment (125ng/µL for Maver-1 and 250 ng/uL for Mino) for another 24h. BCR signaling activity was assessed by measuring phosphorylation levels Syk, BLNK, and Btk using WB. **(f)** Schematic representation depicting SHP-1 mediated regulation of BCR signaling by p53.

To elucidate the mechanisms underlying the repression of BCR signaling mediated by TP53, we performed an integrated analysis of the RNA-Seq and CUT&RUN data, in which *PTPN6* (SHP-1) emerged as a direct *TP53* target (**Fig. 7b, Supplementary Table 21 & 22**). We then examined the protein level of SHP-1 in the Z-138 isogenic pairs and observed a notable increase of SHP-1 expression in the WT cells following p53 activation, whereas the expression levels in the *TP53*-deficient cells remained unchanged. Consistently, in cells with ectopic p53 expression, SHP-1 expression was also significantly increased upon *TP53* induction (**Fig. 7c, d**). To validate this association, we employed siRNA to knock down SHP-1 and identified a combination of three siRNAs that exhibited a superior knockdown effect, which was utilized in the subsequent experiments **(Supplementary Fig. 12c)**. Remarkably, inhibition of SHP-1 restored BCR signaling in cells overexpressing WT-*TP53*, reinforcing the role of SHP-1 as a key mediator of p53-driven BCR signaling suppression **(Fig. 7e)**. Taken together, these findings highlight the important role of the p53-SHP-1 axis in controlling BCR signaling, a mechanism that is disrupted in *TP53*-deficient MCL, potentially influencing disease progression and therapy resistance **(Fig. 7f).**

## Discussion

The pathological and clinical heterogeneity in MCL can be addressed with an in-depth understanding of pathobiology and the evolution of biology-driven classification. Previous research has determined cell proliferation as a critical factor influencing survival outcomes^6^. This proliferation can be effectively quantified at the mRNA level, providing substantial discrimination for clinical application^11,29^. Tumor proliferation is driven by oncogenic activities such as the constitutive expression of CCND1^2,15,16^, and defects in DDR mechanisms, primarily due to somatic inactivation of *TP53*^5,14^. Other factors, encompassing increased genomic complexity^19,20^, alterations in *CDKN2A* (p14/p16) and *TP53*^21,22^, and nodal TME dependent or independent activation of NF-κB signaling^30^ also play a significant role in prognosis, independent of cell proliferation. Moreover, SOX11 overexpression plays a key role in the pathogenesis of MCL by blocking the B-cell differentiation and prompting interactions with TME, although the prognostic significance of SOX11 is controversial^31–34^.

The current study provides a logical extension to earlier studies^2,6,8,12,16^ and has revealed genomic and TME heterogeneity associated with prognostic subgroups. Genetic analyses uncovered significant alterations in pathways related to DDR chromatin reorganization, cell cycle, BCR signaling, and apoptosis. The mutation landscape aligns with previous reports^2,6,8,12,16^, indicating a low tumor mutational burden (∼1 mutation per megabase) but a highly complex CNA profile. The major strength of this study lies in its substantial cohort size, which not only confirms earlier findings^10,12,35–37^ but also establishes the legitimacy of rare variants identified in singular studies. Furthermore, this research sheds light on the impact of specific genetic variants on patient outcomes, particularly in the context of integrating immunotherapy into treatment regimens. We confirmed the mutual exclusivity of *ATM* and *TP53* mutations, as reported in prior research^38,13^. As two of the most frequently altered genes in MCL, especially considering both are critically involved in the DDR mechanism, this exclusivity underscores the importance of DDR defects in the pathogenesis of MCL. Importantly, our study also suggests that while immunotherapy showed improved outcomes in cases with *ATM* mutations, this benefit was not observed in cases with *TP53* or *CCND1* mutations. This underscores the distinct biological and clinical implications of *ATM* and *TP53* mutations in therapy responses. *CCND1*^Y44D^ hotspot mutation, which was reported to increase the protein expression and mediate resistance to ibrutinib^39^, was also identified in this study. However, mutations in the 3’UTR of *CCND1* were only detected using a custom panel, not in the WES cohort, due to a lack of targeted capture regions. Previous studies have shown that alterations in the 3’UTR, such as truncations or mutations, lead to enhanced *CCND1* mRNA stability and are associated with worse clinical outcomes in MCL^40,41^. These findings highlight the intricate involvement of *CCND1* in MCL beyond gene rearrangements. *KMT2C*, *PCLO, BCOR, NOTCH1, NOTCH2* mutations showed lower detection in Coh-1 and higher in Coh-2. Among them, *KMT2C* and *NOTCH1/NOTCH2* mutations are subclonal^13^, which may result in their lower detection in WES (108x depth) compared to TS (500x depth). We also incorporated interaction terms in multivariate modeling to assess the interplay between genetic events and clinical features. *TP53* mutations were associated with poor outcomes regardless of treatment, but their adverse effect was further amplified in patients presenting with B symptoms. This observation reinforces the robustness of our recently established risk assessment model that incorporates both p53 positivity and the presence of B symptoms^16^. Likewise, *CCND1* mutations showed strong interactions with age and B symptoms, underscoring their prognostic significance. These findings highlight the potential utility of a composite prognostic model that integrates both clinical, pathological, and genetic parameters to improve risk stratification and guide more personalized clinical management.

While the genetic subclassification has shed insight on biological diversity or clinical heterogeneity in Diffuse large B cell lymphoma (DLBCL) but is currently evolving in MCL. The prognostic assessment utilizes transcriptomics specifically targeting proliferation markers, such as MCL35^6,42^ and Ki67 index^43^, alongside clinicopathological paratmeters^16^, with a recent proposal for a genetic subclassification^13^. We refined this classification and uncovered six genetic subtypes of MCL with prognostic significance, echoing the study that delineated four genetic subtypes^13^. Consistently across studies, mut-*ATM*/del11q and mut-*TP53*/del17p, emerge as determinants in subgroup differentiation, which highly suggests the critical roles of *ATM* and *TP53* in MCL pathogenesis, leading to tumors with disparate biological characteristics. Beyond these, the remaining four genetic subtypes in our classification inform distinct biological associations, including: mutations in epigenetic regulators (e.g., *KMT2D, KMT2C, SPEN, ARID1A*); loss of tumor suppressors (e.g., *PRDM1*, *del6q*); anti-apoptotic alterations (e.g., *BCL2L10*, *CCND1* mutation, *gain-15q/17q*); and dysregulated differentiation pathways (e.g., *BCL6*, *ATR gain-3q*). Together, these subtypes highlight a complex genomic landscape beyond the canonical *CCND1* rearrangement and emphasize the molecular heterogeneity underlying MCL development. While these subtypes offer valuable biological insight, further studies are warranted to validate their clinical relevance and fully define their implications for tumor behavior. Meanwhile, a simplified classification of MCL cases into low-risk (intact *TP53*, monoallelic *ATM* mutations), intermediate-risk (*ATM* inactivation with 11q loss and epigenetic mutations), and high-risk (biallelic *TP53* inactivation with complex co-deletions) may provide a more practical framework for clinical application.

Unlike other NHLs, the TME features in MCL have not been well characterized. Our single-cell IMC analysis revealed that CD8⁺ T cells constitute the most abundant immune population (11.7%); however, they exhibit high expression of immune checkpoint molecules, suggesting a functionally exhausted phenotype. This raises the possibility that checkpoint inhibition could reinvigorate CD8⁺ T cell activity as a therapeutic strategy in MCL, particularly in light of recent transcriptomic deconvolution data showing that immune-depleted TMEs are associated with poor outcomes and resistance to BTK inhibitors in MCL^44^. Nevertheless, a phase I clinical trial of Nivolumab in relapsed/refractory NHLs included four MCL patients, none of whom responded to the treatment^45^. This may suggest that T cell reactivation alone is insufficient for effective tumor control and that additional immunomodulatory strategies are required. Interestingly, our analysis also found that CD163^+^ myeloid cells, particularly intermediately differentiated CD68⁻ subsets, are abundant in the MCL TME, indicating a skewing toward an M2-like (pro-tumor) phenotype. These findings suggest that tumor-associated macrophages (TAMs) likely contribute to immune dysfunction in MCL. Preventing M2-like (pro-tumor) polarization and promoting M1-like (anti-tumor) differentiation may help restore effective immune surveillance. This strategy could be especially relevant in *TP53*-perturbed MCL, as our data showed a significant increase in spatial neighborhoods characterized by interactions among SOX11⁺ tumor cells, CD68⁺/⁻ myeloid cells, and CD8⁺ T cells. We note inclusion of more IMC-characterized cases would further support these findings.

Additionally, the *TP53*-perturbed group exhibited a distinct immune signature marked by increased expression of CXCR3, cMYC, pNF-κB, and immune checkpoint molecules including LAG-3 and TIM-3, indicating a dynamic interplay between immune activation and suppression. One plausible mechanism is that *TP53* loss enhances NF-κB signaling, a central regulator of inflammation and immunity, leading to increased immune cell recruitment via CXCR3 and HLA-ABC upregulation. However, NF-κB activation also drives expression of checkpoint molecules such as LAG-3 and TIM-3, promoting T cell dysfunction and immune escape. In parallel, the observed upregulation of cMYC suggests a shift toward metabolic reprogramming, which may further support immune resistance and tumor progression. Thus, *TP53* perturbation appears to generate a highly inflammatory yet immunosuppressive microenvironment, where immune infiltration is robust but cytotoxic function is impaired. These findings align with recent studies indicating that *TP53* loss reshapes the tumor-immune landscape, enabling immune escape^46,47^. Proposed underlying mechanisms include increased cytokine secretion, enhanced integrin recycling, and the transfer of mutant *TP53* via exosomes to surrounding cells within the TME^46–48^. Report of limited PD-1 expression in MCL highlight the need to explore alternative immune checkpoints^49^. In *TP53*-perturbed cases, combining these targets with NF-κB or metabolic inhibitors may more effectively restore anti-tumor immunity.

As a well-established tumor suppressor, *TP53* has been thoroughly examined for its target genes and functions^50,51^. However, there is a significant knowledge gap, particularly regarding specific disease context. Our investigation not only corroborated the established role of p53 in regulating cell cycle and apoptosis but also unveiled its role in regulating BCR signaling. This is in accordance with the clinical observation indicating limited efficacy of the BTK inhibitor ibrutinib in patients with *TP53* mutations^26^. We revealed that the activation of *PTPN6 (SHP-1)*, a key inhibitor of BCR signaling, plays a significant role in repressing BCR signaling, which potentiates a strategy of modulating SHP-1 for enhancing the efficacy of ibrutinib in MCL with p53 alterations. Additionally, previous studies have demonstrated that BTK mutation-mediated ibrutinib resistance is rare in MCL, whereas adaptive reconfiguration of the kinase signaling network, especially the PI3K-AKT and β1-integrin pathways, has a pivot role in developing resistance to ibrutinib^52–54^. Therefore, our findings also raise the intriguing possibility that SHP-1 deactivation may facilitate kinase pathway rewiring and contribute to ibrutinib resistance in MCL, even in the absence of *TP53* deficiency, given that it is a major phosphatase negatively regulating multiple kinase pathways. Future study is needed to elucidate these mechanisms, paving the way for improved therapeutic strategies for MCL.

## Data availability

**Raw Data dbGaP Study Accession**: phs003849.v1.p1. All raw data related to RNA-Seq/CUT&RUN assay has been deposited to NCBI GEO with respective GEO ID’s: Coh-1 RNA-Seq (GSE271664), CUT&RUN MCL Cell lines (GSE271594), RNA-Seq MCL Cell lines (GSE271503).

## Processed Data

All processed data has been added as **Supplementary Tables** for Genomics, Tumor Microenvironment, and CUT&RUN assays along the main text. Sequencing data for Cohorts 1 and 2 are deposited in dbGaP (phs003849.v1.p1). RNA-Seq data for Cohort 1 is in NCBI GEO (GSE271664). RNA-Seq and CUT&RUN data for MCL cell lines are under GSE271503 and GSE271594, respectively. Additional data generated in this study are provided in the Supplementary tables.

## Code availability

The original code related to the genomics/Tumor microenvironment/RNA-Seq/CUT&RUN analysis is available on **GitHub** (https://github.com/sharmas30-94/MCL_2024).

## Methods

### Patient specimens

The study analyzed 290 MCLs, consisting of two study cohorts: Cohort 1 (Coh-1, 130 cases from the post-rituximab era: NAMCLP + 23 cases from a previously published study^37^) and Cohort 2 (Coh-2, 137 cases from the pre-rituximab era). In Coh-1, 15 were rituximab-naïve with 5 receiving hyper-CVAD therapy and 10 undergoing other treatments. Coh-2 included 36 rituximab-treated and 57 rituximab-naive cases. Coh-1 was subjected to Whole Exome Sequencing (WES) with an average depth of 108x, while Coh-2 underwent targeted sequencing with average depth of 557x, employing a custom gene panel targeting genes frequently mutated in NHL as established in prior studies^17^. For somatic variant detection, diagnostic biopsy DNA was used, and normal tonsil DNA was employed to identify potential germline variants in SCNA analysis. Expression analysis for Coh-1 utilized RNA-Seq data from 47 samples, while Coh-2 analysis was based on microarray data (HG-U133plus2 platform) of 25 cases^8^. To confirm the reproducibility of mutation identification, 17 MCLs from Coh-1 were sequenced using both WES and TS. The study received approval from the Institutional Review Boards of all involved institutions.

The median age at diagnosis was 61 years in Coh-1 and 62 years in Coh-2. Most cases in both cohorts were diagnosed at late stages (III & IV), accounting for 89.4% (110/123) in Coh-1 and 87.3% (62/71) in Coh-2. In Coh-1, 55.8% (82/147) of patients were aged ≥60 years, compared to 57.3% (59/103) in Coh-2. SOX11 positivity was observed in 88.0% (95/108) of tested cases in Coh-1, while Ki-67 positivity was found in 29.8% (31/104). Rituximab was administered as part of the primary regimen to 87.1% (128/147) of patients in Coh-1 and 36.8% (28/76) in Coh-2, with 12.9% (19/147) and 63.2% (48/76), respectively, being Rituximab-naïve. The MIPI scores in Coh-1 indicated that 43.8% (21/48) of patients had a low score, 45.8% (22/48) had an intermediate score, and 10.4% (5/48) had a high score. Male patients comprised 49.0% (72/147) in Coh-1 and 83.1% (64/77) in Coh-2. Classical pathological subtypes were observed in 82.1% (55/67) of patients in Coh-1, while 17.9% (12/67) had blastoid/pleomorphic subtypes. A diffuse growth pattern was noted in 29.4% (20/68) of Coh-1, while the remaining cases exhibited other growth patterns. Regarding race, 94.2% (113/120) of Coh-1 were Caucasian, while 5.8% (7/120) identified as other races. Disease site analysis in Coh-1 revealed that 18.7% (23/123) of patients had nodal disease, 10.6% (13/123) had extra-nodal disease, and 70.7% (87/123) had both nodal and extra-nodal involvement. B symptoms were present in 28.5% (35/123) of Coh-1, while 71.5% (88/123) did not report B symptoms. Additionally, 34.3% (48/140) of cases in Coh-1 underwent hematopoietic stem cell transplantation (HSCT), while 65.7% (92/140) did not.

### DNA/RNA isolation and library preparation for high-throughput analysis

Total RNA and g-DNA from formalin-fixed-paraffin-embedded (FFPE) were extracted using RNAstorm^TM^/DNAStorm^TM^ Kit (Celldata) as per manufacturer’s guidelines (Catalog Number: CD506, CD507). The quality and quantity of RNA/DNA were assessed using Qubit fluorometric quantification and an Agilent Bioanalyzer. Genomic DNA was sheared to 250 bp, underwent end-repair, A-tailing, adapter ligation with Illumina paired-end adapters, and amplification. The samples were column purified, quantified, and hybridized overnight to DNA baits from the SureSelect Human All Exon V7 kit (Agilent Technologies). The enriched libraries were further amplified and sequenced on an Illumina HiSeq 2500. For RNA sequencing (RNA-Seq), samples with RNA quality of >35% greater than 200 bases were used. A minimum of 200 ng of total RNA was converted into mRNA libraries using the KAPA RNA HyperPrep with RiboErase, followed by sequencing on the NovaSeq 6000 platform.

### Sequencing analysis

We conducted WES on 130 cases from Coh-1 and TS on 137 cases from Coh-2 using a custom-lymphoma panel of 380 genes^17^. Variant calling was performed using the same pipeline for both cohorts, with the only difference being a lower Variant Allele Frequency (VAF) cutoff (3%) for TS^55^ compared to WES (5%)^56^. The quality of the raw reads was assessed by FastQC (v 0.11.7), and adapter sequences along with poor-quality WES reads were subject to trimming with Trimmomatic (v0.36)^57^. The processed reads were aligned to the human genome (hg38) with BWA (0.7.17-r1188), and duplicate reads were marked using Picard (v2.9.0) (https://broadinstitute.github.io/picard/). Variants were annotated with Annovar and retained if they met specific criteria: at least four reads in the tumor sample and with variant reads present on both the plus and minus strand^58^. Somatic mutations were identified using a combination of Varscan2 *and* Mutect2 (v4.1.8.1)^59–61^. To ensure high-confidence calls, variants were included only if they met at least one of the following four criteria: (1) sites not flagged in dbSNP (i.e., marked as “.”), called by Varscan2 (Y), and also detected by Mutect2; (2) TP53 mutations unflagged in dbSNP, called by Varscan2, and detected by Mutect2; (3) sites unflagged in dbSNP, called by Mutect2, detected by Varscan2, and marked as “PASS” in the VCFFILTER.Mutect2 field; (4) sites with population allele frequency (WES.non_cancer_AF_popmax) < 1%, and VAF > 5% in both Mutect2 and Varscan2 or > 3% for TS. Additionally, mutations in genes previously reported to be highly and recurrently mutated across multiple cancer types often due to sequencing artifacts or hypermutability were excluded based on established blacklists^62^. The selected SNV data was analyzed using oncoplot from maftools R package^63^. The 23 external cases included in Coh-1 were processed using a different pipeline^37^; however, integration was performed at the variant level, not at the pipeline level.

The RNA-seq analysis was performed on 47 MCL samples from Coh-1, derived from FFPE tissues. Raw read quality was assessed with FastQC (v 0.11.7) and sequences from contaminating adapters were removed with Trimmomatic (v0.36)^64^. Trimmed reads were then aligned to the hg38 genome using STAR (2.7.10b)^65^. RNA-Seq levels were quantified with HTSeq (v0.9.1)^66^ and normalized using Trimmed Mean of M-values (TMM) in edgeR^67^ for GSEA and iGC input. Functional annotation of differential genes was performed using ConsensusPathDb^68,69^. The microarray gene expression data (Coh-2, n=25) was normalized using the limma R package^70^. We used CIBERSORTx^71^ with LM22 and 1000 permutations on TMM-normalized counts.

### Clinical outcome correlation and statistical analyses

The Kaplan-Meier method was used to estimate the overall survival distributions using the R survminer package (https://github.com/kassambara/survminer). Overall survival was defined as the period from diagnosis to either the date of death or last contact, with patients alive at the last contact considered censored. The log-rank test was used to compare survival distributions^72^. All statistical tests are two-sided, and p-values < 0.05 were considered statistically significant unless otherwise specified. Multivariate analysis employed Cox Hazard ratio models with significance set at p-values < 0.05. Data analysis was performed using R (R Project), and visualizations were created with ggplot2 and survival R packages.

### Somatic copy number aberration analysis (SCNA)

CopyWriteR was utilized to generate genome-wide copy number profiles from BAM files that had been mapped, realigned, sorted, and deduplicated, enabling the identification of off-target regions^73^. A bin size of 100kb and hg38 assembly was used to perform the segmentation using the circular binary segmentation (CBS) within the DNAcopy Bioconductor package after the correction of GC-content and removal of blacklisted regions. Of note, for 130 MCLs from Coh-1, we used a normal tonsil sample as a polymorphism control. For classifying CN variations from genomic data, we defined four categories based on the raw mean copy number intensity of each segment. Segments with an intensity >= log base 2 of 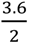 were classified as CN amplifications (Absolute Copy = “4”), indicating significant copy number increases. Those with intensities >= log base 2 of 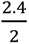 but below the amplification threshold were marked as CN gains (Absolute Copy = “3”), representing moderate copy number increases. Segments at or below log base 2 of 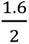 were categorized as CN losses (Absolute Copy = “1”), denoting notable reductions in copy number. Finally, segments at or below log base 2 of 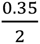 were considered as homozygous losses (Absolute Copy = “0”). The percent aberrant genome per case was calculated by using CNAapp^74^. We obtained segmented data on 23 MCLs included in Coh-1 from a previously published series^37^ and merged with 130 MCLs by filling in the missing segments to account for the consistency of segment size. For Coh-2, no sample controls were used. Segmented data were analyzed using GISTIC2.0^18^ to identify recurrent CNAs within the GenePattern (https://www.genepattern.org/) with default parameters and reported as copy loss or gain if the absolute DNA copy number was =< 1.6 or >=2.4, respectively. Literature and adjusted p-value filtering ensured robust results. SCNA driver alterations, significant at 90% confidence interval, were identified. CNAapp predicted focal, global, and broad-level copy number burden^74^. GenVisR and gisticChromPlot (maftools) facilitated frequency plot visualizations^63,75^. Integrative GEP and SCNA analysis was performed using iGC^19^ R package.

### Somatic interaction analysis

To evaluate the mutual exclusivity and co-occurrence of genomic events in MCL, we combined the two cohorts based on common mutations and common copy number aberrations and applied the somaticInteractions function using maftools^63^.

### NMF consensus clustering analysis

To define molecular subtypes associated with clinical outcomes, we began with a curated list of 51 significant genomic events identified across the Coh-1, including recurrent mutations, copy number gains, and deletions. Initial unsupervised clustering using the full feature 51 set did not yield robust prognostic stratification. To refine feature selection, we implemented a randomized feature subset approach, generating 100 random sets, each containing 35 genomic features. For each feature set, non-negative matrix factorization (NMF) consensus clustering was performed using the NMF consensus clustering^76^, exploring a range of cluster numbers (K = 4 to K = 7) for each feature set. Clustering results were evaluated using two primary criteria: (1) the emergence of a distinct cluster enriched for biallelic *TP53* and *ATM* alterations, reflecting known high-risk biology, and (2) evidence of prognostic separation in at least two out of the three K values, based on log-rank survival analysis. Based on these criteria, feature set 86 was selected for downstream analysis. To identify genomic events uniquely enriched within each cluster, we performed Fisher’s exact test, comparing the frequency of each genetic event across clusters. A threshold of p < 0.1 was used to define statistically enriched events. The resulting binary matrix of genomic alterations (mutations, amplifications, deletions) was visualized using the ComplexHeatmap^77^ package in R, generating an oncoplot that highlights the genomic architecture of each cluster.

### Random Forest classification

To predict molecular clusters in a second cohort (Coh-2) based on targeted sequencing features, we trained a supervised classification model using WES data from Coh-1, where NMF-based cluster assignments were previously established. Genomic features from Coh-1 included binary indicators (presence/absence) of gene-level deletions, gains, and mutations. To ensure consistent input for both training and testing, we identified the intersection of features present in both Coh-1 and Coh-2 and trained the model using only this shared set of alterations. A Random Forest classifier was implemented using the randomForest package in R. The model was trained on Coh-1 using 500 trees (ntree = 500), with default settings for mtry (number of variables randomly sampled at each split) and nodesize (minimum size of terminal nodes). No imputation or class rebalancing was applied. The trained model was used to assign cluster labels in Coh-2 based on their genomic profiles. Model performance was evaluated by comparing the predicted labels to original NMF-assigned clusters using the caret package. Evaluation metrics included overall accuracy and per-class precision, recall (sensitivity), F1-score, and support (sample count per cluster) (**Supplementary Table 15**). This feature-intersection approach ensured that the classifier was trained and tested on comparable input, minimizing bias introduced by missing data or artificial imputation.

### Samples included in IMC analysis

The dataset consisted of 50 samples, including 42 sequenced samples from Coh-1 (n=28) and Coh-2 (n=14), 5 unsequenced NAMCLP cohort cases from the original study with available paraffin blocks, and 3 controls (spleen, tonsil, and lymph node). The 42 sequenced cases were classified into three genomic groups based on *ATM* and *TP53* alterations: *ATM*-perturbed, defined as cases with either an *ATM* mutation or *ATM* copy number loss, or both; *TP53*-perturbed, defined as cases with either a *TP53* mutation or *TP53* copy number loss, or both; and WT (wild-type), defined as cases with no detectable alterations in *ATM* or *TP53*. The 5 additional unsequenced samples were included to improve immune phenotyping and, where material was available. To ensure adequate representation of the *TP53*-perturbed group, we included single-slide cases specifically for this group. In total, nine tissue slides were analyzed using the Hyperion platform, consisting of seven unstained single-tissue slides (all *TP53*-perturbed) and two Tissue Microarrays (TMAs) slides. All tissue cores and slides were assessed by two pathologists, who identified Regions of Interest (ROIs) on H&E-stained slides for downstream analysis. Due to tissue block availability limitations, whole slides were included, but only the identified ROIs on those slides were used for our analysis. A summary distribution of the 50 samples across the two cohorts, TMA origin, number of ROIs ablated, and averaged phenotype and spatial fractions per sample has been detailed (**Supplementary Table 18**). Notably, at least one ROI was ablated for all 50 samples, ensuring their inclusion in downstream analyses. The selection of samples was based entirely on the material availability.

### Tissue microarray

The TMAs were sectioned at 4-µm thickness, baked at 60°C for 90 minutes, de-waxed in xylene, and rehydrated through a graded alcohol series. Antigen retrieval was carried out in Tris-EDTA buffer at pH 9 at 95°C for 30 minutes, followed by blocking with 3% BSA in PBS for 45 minutes, and overnight incubation at 4°C with a 36-antibody cocktail tagged with rare lanthanide isotopes. The slides were analyzed using the Fluidigm Hyperion Tissue Imager, which combines laser ablation with mass spectrometry^78^. A 36-biomarker panel was generated to define the canonical immune subsets and functional state markers based on our literature review. Each of the biomarkers were thoroughly validated through literature review, vendor validations, and rounds of staining validations consisting of using appropriate tissues for each biomarker and titration level determination by serial dilution (1:100, 1:400, 1:800, 1:1000). Laser beam of 1-µm^2^ spot size ablated tissue areas of 1000-µm^2^ per core at a frequency 200 Hz, measuring metal isotopes and mapping them against each spot to create intensity profiles and digital spatial maps of the tissues. Detailed descriptions of the ablation techniques have been previously described^79,80^. Raw IMC data was acquired and archived in the MiniCAD Design File (MCD) format.

### Image segmentation

Image segmentation utilized the DeepCell Mesmer model^81^ (version 0.11) with standard parameters, a deep learning framework trained on various multiplex imaging modalities, including IMC. We used the arcsinh-transformed nuclear channel (191Ir) image, scaling pixels to the range [0, 1], and the arcsinh-transformed HLA-ABC counts image as a membrane channel. The final membrane channel was the result of subtracting the gaussian filter of the image at sigma=3 from the gaussian filter with sigma=1 and again scaling to unit range. To obtain our single-cell dataset, we averaged each marker’s expression over pixels belonging to each cell. To address technical variance from tissue preservation, staining, and ablation in the absence of exact replicates we leveraged established batch normalization techniques. Given that IMC is a single-cell resolution platform and expression measurements are unbiased with respect to cell size, we judged cyCombine^82^, which employs an empirical Bayesian method to reduce the Earth Mover’s distance among batch distributions for each marker to be the method of choice at the time of analysis, due to it being a platform-agnostic method and with the advantage that it is replicate-free and adaptive to the noise profile that best correlates with the observed batch effect. It performs cell expression corrections via using an empirical Bayesian approach to minimize the earth mover’s distance between similar clusters across different batches.

### Phenotypic clustering

The dimensionality of highly multiplexed data requires an automated and unsupervised approach to phenotyping in the form of clustering. We first normalize cell expression by clipping at an elbow point at the upper end of expression among cells in each marker (outliers can be present for a range of reasons, including hot pixels and local segmentation errors), and then scaling all markers such that the range is between 0 and 1. Using the normalized dataset, hierarchical Ward-based clustering was applied on lineage markers to a set number of clusters within each ROI, followed by a meta-clustering across all ROIs employing the Leiden community clustering method^83,84^. Using heatmaps of mean expression per cluster, we manually merge and annotate clusters into the final phenotypic labels relying on expected knowledge, such that each cell in the dataset is assigned a phenotypic label (e.g., CD68^+^ Myeloid, CD4^+^ T cell). Single-cell data were phenotyped using a lineage and MCL-relevant subset of the IMC panel, including CCND1, SOX11, canonical markers of B, T, and myeloid cells. Hierarchical clustering was applied to all 106 ROIs individually using the Ward method with Euclidean distances as implemented in the fastcluster^85^ and scikit-learn Python package, obtaining 128 clusters per ROI. These were then meta-clustered via Leiden^86^ into 32 clusters. The meta-clusters were validated and annotated by an expert hematopathologist, and clusters representing the same phenotype were merged. Our final phenotyping serving as the basis of subsequent analysis, identified all expected major subsets, including CD4^+^ T, CD8^+^ T, Tregs, myeloid, endothelial, and neoplastic SOX11^+^ and SOX11^-^ cells, delineating 9 phenotypes in total.

### Spatial analysis

To characterize the interactions of cells, we use their locations and labels to infer spatial motifs representing modes of local phenotypic milieus and one or more scales which are defined by setting an interaction relevance score based on distance between cells. Using 25- and 50-micron (equivalent to pixel in IMC) radii, for any given cell which we term the source cell, we assign a score of interaction to neighboring cells which is inversely proportional to the distance between the pair’s centroids and falls off to zero beyond said radius. By averaging interactions per phenotype with the source cell contributing, we assign a cell to phenotype interaction score for the entire dataset. We leverage the fact that this resulting dataset has a form analogous to the expression dataset to apply the same clustering techniques as used for phenotyping towards identifying spatial motifs, or neighborhoods, which here we use interchangeably. By quantifying the average functional marker expression of phenotypes, their abundance fractions, and those of 25 micron-, and 50 micron-scale neighborhoods in each ROI and averaging ROIs when more than one is available per patient, we obtain a combined feature set to use in identifying spatially informed microenvironment-based predictors of clinical variables.

### Functional/state marker expression differences on phenotypes across genetic subgroups

To assess differences in immune functional states across genetic subgroups, we analyzed marker expression across immune phenotypes using spatially resolved transcriptomic or single-cell imaging data. The input matrix consisted of annotated ROIs per patient, where each row corresponded to a unique combination of ROI (txt), and immune phenotype (e.g., CD8⁺ T cells, Tregs, DCs, macrophages). Each column captured the average expression of a functional or lineage-specific marker (e.g., PD-1, CD20, FOXP3, CXCR5, LAG-3, CD3, CD45RA, CD4, CD11b) within that phenotype-region pair. We focused on nine major immune phenotypes identified within the TME, and summarized expression by computing the average marker intensity per phenotype per patient. This was done to reduce intra-sample variability and facilitate comparisons across genetically defined subgroups. Each patient was assigned to a genetic subgroup based on somatic alteration profiles (e.g., *ATM*-perturbed, *TP53*-perturbed). Marker expression values were compared between genetic subgroups within each immune phenotype using the. Statistical comparisons were performed using the Kruskal-Wallis test (p < 0.05).

### Cell line construction and gene knockdown

Three MCL cell lines, Maver-1, Mino, and Z-138 were utilized in this study. All cell lines were cultured in RPMI-1640 medium supplemented with 10% Fetal Bovine Serum (FBS) and 1% Penicillin/streptomycin. To generate inducible p53 expression in Maver-1 and Mino cells, wild-type (WT)-*TP53* was cloned into a lentiviral Tet-On 3G inducible expression vector (Tet-pLV-TP53). For lentivirus production, 293T cells were transfected with Tet-pLV-TP53p, CMV-dR8.2 dvpr, pMD2.G using PureFection™ transfection reagent (System Biosciences, Palo Alto, CA, USA). Viral supernatants were collected 48h and 72h post-transfection and centrifugated at 1000 rpm to remove cell debris. The supernatants were then filtered through 0.45 mM filters and concentrated overnight at 4°C using a lentiviral concentrator (Origene, Rockville, MD, USA). Mino and Maver-1 cells were then transduced with the concentrated lentivirus using TransDux^TM^ Max transduction reagent (System Biosciences) and incubated for 48h, followed by puromycin selection. Single clone selection was then performed and the p53 expression was evaluated by western blotting (WB) after 24h of Doxycycline (Dox, Sigma-Aldrich, St. Louis, MO, USA) treatment. To create *TP53* Knock-out (KO) or mutation in WT Z-138 cells, various all-in-one CRISPR constructs were developed. Specifically, fidelity SpCas9^23^, CEB4max-SpG^25^, and AncBE4max^24^, were used to create p53-KO, -R248Q, and -R273C, respectively. Z-138 cells were electroporated with the CRISPR vectors using the Neon transfection system (Thermo Fisher Scientific, Waltham, MA, USA). GFP-positive cells were sorted by flow, followed by single clone selection. Sanger sequencing was performed to identify clones with correct editing. For the *PTPN6* knockdown, Z138, Mino, and Maver-1 isogenic cells were transfected with siRNA mixtures or negative control (NC1) (IDT, Coralville, IA, USA) using Amaxa 4D nucleofector (Program CM-116 for Z138 and Program CM-150 for Mino and Maver-1). The sequences of siRNAs used are listed in **Supplementary Table 20.**

### RNA-Seq analysis

For Z138, Mino, and Maver-1 isogenic cell lines, raw sequencing data were processed using a standardized RNA-seq pipeline. Initial quality assessment of raw reads was performed using FastQC (v0.11.7). Adapter sequences were removed using Trimmomatic (v0.38), and the resulting trimmed reads were aligned to the human reference genome (hg38) using STAR. Duplicate reads were marked using Picard (v2.9.0), and SAMtools (v1.3.1) was used to remove unmapped and non-primary aligned reads from the resulting BAM files. HTSeq-count was used to generate raw gene-level count matrices. These counts were then used as input for differential expression analysis using DESeq2 (via GenePattern with default settings). For the Mino and Maver-1 lines, samples harboring p53 mutations were grouped and compared against their respective p53-CTRL (control) samples. Similarly, for Z138, samples modified with mutant *TP53* alleles (R273C and R248Q) were analyzed together and compared to WT-*TP53* samples. Differential expression was assessed using a cutoff of log₂FoldChange ≥ 2, p-value < 0.05, and adjusted p-value (padj) < 0.05 to determine statistically significant gene expression changes.

### CUT&RUN assay

CUT&RUN assay was performed on Z-138, Mino, and Maver-1 isogenic cells using the CUTANA^TM^ ChiC/CUT&RUN Kit (Epicypher, Durham, NC, USA) following the manufacturer’s instructions with p53 antibody (polyclonal, Epicypher #13-2015) used at 0.5 µg per reaction and 2 replicates per experimental condition were used. Data analysis was executed following previously established methods^87^. Raw read quality was assessed by FastQC (v 0.11.7), adapters trimmed (Trimmomatic, v0.38) and trimmed reads aligned to hg38 and to the spike-in reference (Escherichia_coli_K_12_MG1655) control with Bowtie2 (2.4.4). Duplicates were marked with Picard (v2.9.0) and unmapped and non-primary aligned reads were removed from bam files (samtools v1.3.1). Bam files were converted to bed file format (bedtools v2.27.1) and filtered to retain read pairs on the same chromosome that have a fragment length <1000 base pairs. Samples were normalized to a scaling factor derived from the reads aligned to the E. coli spike (https://yezhengstat.github.io/CUTTag_tutorial/index.html [yezhengstat.github.io]). Peaks were called with SEACR (v1.3)^88^. Seacr peaks were merged for all p53 cut and run samples using GRanges R package and then a count matrix for the peak regions and the samples was created^89^. Differential peaks were identified using DESeq2, and promoter regions with log₂ fold change ≥ 1 and p-value < 0.1 were selected for visualization. deepTools (v3.5.1) was used to create the bigwig for peak visualization^90^. Motif analysis was performed using HOMER (v4.11.1)^91^. Peaks from SEACR were first merged across replicates using mergePeaks, retaining only regions shared across replicates. The resulting consensus peak sets were then compared to the IgG control peaks using mergePeaks, and peaks overlapping with IgG were excluded. HOMER’s findMotifsGenome.pl was run on the filtered consensus peaks using default parameters to identify top 10 enriched sequence motifs. IGV genome browser was used for peak visualization^92^.

