## Supplementary Text and Figures for "Functional genomics and tumor microenvironment analysis reveal prognostic biological subtypes in Mantle cell lymphoma"

#### Supplementary Methods

##### RNA-seq and Gene Set Enrichment Analysis (GSEA)

RNA-seq data were available for 46 cases from Coh-1, for which FPKM expression counts were generated. These counts were normalized using TMM (Trimmed Mean of M-values) normalization as implemented in the edgeR package to account for compositional differences across samples. Normalized expression data were then used as input for Gene Set Enrichment Analysis (GSEA), performed using the GSEA module from GenePattern (https://www.genepattern.org/) with default parameters. GSEA was initially performed using the MSigDB Hallmark and WikiPathways databases to identify biologically relevant pathways associated with risk stratification. For each group comparison: Low-risk vs. others, Intermediate-risk vs. others, and High-risk vs. Others. GSEA was used to detect pathways significantly enriched in each risk category. Subsequently, pre-ranked GSEA was conducted using gene lists ranked by differential expression metrics to refine the analysis and enhance sensitivity to pathway-level changes. Gene sets showing significant enrichment were further interrogated by extracting their leading-edge genes, representing the core subset of genes driving the enrichment signal. To visualize and compare pathway activity across the three groups, normalized expression values of the leading-edge genes were averaged per pathway within each risk group, enabling the characterization of pathway-level expression dynamics in relation to risk status.

##### Cox Proportional Hazards Analysis of Clinical Parameters

Univariable and multivariable Cox proportional hazards regression models were employed to assess the association between clinical parameters and both overall survival (OS) and progression-free survival (PFS). Results are reported as hazard ratios (HRs) with 95% confidence intervals (CIs) and corresponding p-values. All analyses were conducted using the survival package in R, with statistical significance defined as $p < 0.05$. To evaluate potential effect modification by mutation status, we incorporated interaction terms between specific gene mutations (e.g., *TP53*, *ATM*) and clinical covariates (e.g., age, transplant status) into multivariable models. The interaction was modeled using the formula: Surv(time, status) ~ mutation + clinical_variable + mutation × clinical_variable. A statistically significant interaction term suggests that the prognostic effect of the clinical variable differs based on mutation status.

**RNA Sequencing of TP53 modified cell lines**

To profile p53 regulated genes in MCL, Z-138 isogenic cell lines were treated with vehicle control or 500nM Amg-232 (MedChemExpress, Monmouth Junction, NJ, USA) for 16h while Maver-1 and Mino isogenic cells were treated with vehicle control or Dox at 125 and 250 ng/ml for 12h. Total RNA was extracted using the Quick-RNA™ Mini prep kit (Zymo Research, Irvine, CA, USA). The RNA quality was assessed with Agilent Bioanalyzer 2100 (Agilent Technologies, Santa Clara, CA, USA), selecting samples with an RNA integrity number (RIN) greater than 8 for library preparation. The libraries were then sequenced on the Illumina NovaSeq 6000 instrument to obtain ~20 million, 150 bp paired end reads. The reads were aligned to hg38 using the STAR aligner (version 2.5.3)[3]. Gene expression levels were quantified using HTSeq (v0.9.1)[4]. The resulting counts were normalized and analyzed with the R package DESeq2[5] to identify differentially expressed genes.

**Western blot (WB)**

Total protein was isolated from cultured cells using M-PER™ Mammalian Protein Extraction Reagent (Thermo Fisher Scientific, Waltham, MA, USA) supplied with Halt™ Protease and Phosphatase Inhibitor Cocktail (Thermo Fisher Scientific, Waltham, MA, USA). The isolated protein was denatured at 70°C for 10 mins using NuPAGE LDS Sample Buffer supplemented with reducing buffer (Thermo Fisher Scientific, Waltham, MA, USA). Protein electrophoresis was performed using Bolt 4~12% Bis-Tris Plus Gels and MES SDS running buffer (Thermo Fisher Scientific, Waltham, MA, USA). Proteins were then transferred onto nitrocellulose membrane, blocked with Odyssey TBS Blocking Buffer (LI-COR, Lincoln, NE, USA), and incubated with primary antibody overnight at 4°C. The membrane was then washed and incubated with fluorescent secondary antibody (LI-COR) for 1h at room temperature. After washing, the membrane was scanned on the Odyssey CLx imager (LI-COR). Protein was quantified based on band intensity using the Image Studio software (LI-COR). Protein expression change was evaluated by normalizing with β-actin or target protein input. The primary antibodies used for WB are listed in **Supplementary Table 21**.

#### Supplementary Figures

**Supplementary Fig.1: Identification of clinical factors associated with patient survival using Cox regression and Kaplan-Meier analysis.**

**(a)** Cox proportional hazards analysis of clinical and molecular parameters in Coh-1. Forest plots summarizing hazard ratios (HRs) and 95% confidence intervals (CIs) for overall survival (OS, red) and progression-free survival (PFS, blue) based on clinical and molecular variables, including SOX11 IHC, TP53 IHC, Ki-67, nodal status, B symptoms, transplant status, age, and rituximab use. Statistically significant associations (green text) were observed for SOX11 IHC, B symptoms, transplant status, age, and rituximab. The vertical dashed line indicates HR = 1; HRs to the right reflect increased risk, while those to the left indicate reduced risk.

**(b)** Kaplan-Meier survival curves for OS based on TP53 IHC and rituximab administration in Coh-1. Risk tables are displayed below each plot, with groups distinguished by color.

**(c)** Kaplan-Meier survival curves for OS and PFS by B symptoms, transplant status, and biological age in Coh-1.

**Supplementary Fig.2: Genomic variant analysis and signature profiling across Coh-1 and Coh-2.**

**(a)** Variant classification and frequency distributions for two cohorts: Coh-1 (n=153) and Coh-2 (n=137). The upper panels display the distribution of variant class, variant type, and SNV class for each cohort. The variant classification summary highlights the overall mutation profile in each cohort. Variant per sample distribution is shown as histograms.

**(b)** Overview of WES and TS platforms used for genomic profiling. The figure illustrates the genomic breadth of WES compared to targeted sequencing, showing that 94% of the targeted variants were recovered in the WES profiles, with only 6% of variants remaining uncovered. The VAF correlation between common variants detected on the TS versus WES is also shown, with a moderate correlation ($r = 0.68$, $p < 2.2e-16$).

**(c)** Single Base Substitution (SBS) and Legacy Signatures of mutations from Coh-1. The left panel shows the cophenetic metric for clustering SBS signatures. The top three most relevant mutation signatures are matched based on cosine similarity to known mutagenic patterns, including SBS5 (deficiency in base excision repair due to inactivating mutations in NTHL1), SBS30 (exposure to tobacco mutagens), and SBS4 (another tobacco-related mutagenic signature). The right panel displays the top matches for legacy mutagenic signatures from COSMIC with the best-matching signatures listed, including exposure to tobacco mutagens and other unknown mutagenic causes. Of note, SBS and legacy mutational signature analysis was not performed on Coh-2, as the targeted panel only covers a limited set of genes preventing the identification of broader mutational patterns as Coh-1.

**Supplementary Fig.3: Mutational landscape in Coh-2.**

**(a)** Overview of somatic mutations in Cohort 2 (n = 137), displaying mutation types and frequencies across selected genes. Mutation types, including missense, frameshift, splice site, and nonsense are shown for each gene. Clinicopathological features are shown in the bottom heatmap including biopsy type, SOX11 and TP53 IHC, Ki-67 index, rituximab administration, biological age, clinical stage, and sex.

**(b)** Comparison of mutation frequencies for top mutated genes between Coh-1 and Coh-2. Stacked bar charts illustrate the distribution of mutation types in each cohort. Differences in gene-level mutation frequencies were evaluated using Chi-square tests.

**Supplementary Fig.4: Cancer hotspot analysis using cBioPortal MutationMapper.**

**(a-c)** Mutation landscape of (a) *TP53*; (b) *ATM*; (c) *CCND1*. The colors of the lollipops represent different types of mutations. Annotations highlight key cancer hotspots and the corresponding amino acid alterations.

**Supplementary Fig.5: Multivariate Cox regression analysis of CNV lesions and clinical factors in Coh-1**. The forest plot shows hazard ratios and 95% confidence intervals for individual CNV lesions and their interaction with clinical variables including age, B symptoms, transplant status, and immunotherapy use. Significant associations ($p < 0.05$) indicate CNV lesions and clinical features that contribute to OS survival risk in MCL patients.

**Supplementary Fig.6: Somatic interaction analysis across 290 patients.** Each cell in the corrplot represents the odds ratio for either the co-occurrence or mutual exclusivity of common mutations and copy number aberrations, calculated using Fisher’s Exact Test. An odds ratio less than 1 indicates mutual exclusivity, while an odds ratio greater than 1 suggests co-occurrence. The color gradient illustrates the magnitude of the odds ratio, with blue denoting mutual exclusivity and red indicating co-occurrence.

**Supplementary Fig.7: Prognostic NMF subtyping and clustering evaluation in Coh-1.**

**(a)** Schematic of the workflow for identifying prognostic subtypes using genomic data. A total of 100 random sets of 35 genomic features were generated and clustered using NMF across K-values from 4 to 10. The optimal feature set and K-value were selected based on their ability to recover known alterations (e.g., *TP53* and *ATM* loss), identify novel subtypes, and stratify prognosis. Fisher's exact test was used to identify enriched or depleted events in each subtype, based on odds ratios and p-values.

**(b)** Cophenetic correlation coefficients for k = 4 to k = 10.

**Supplementary Fig.8: Copy number aberration burden score and immune landscape across merged clusters.**

**(a)** Violin plots showing the distribution of FCS, BCS, and GCS scores across merged molecular clusters stratified by OS. Patients were grouped into low-risk (black), intermediate-risk (gray), and Poor OS (red) clusters. Boxplots are overlaid to indicate the median and interquartile range.

**(b)** Boxplots showing the distribution of inferred immune cell proportions across OS-based merged clusters, based on CIBERSORTx deconvolution.

**Supplementary Fig.9: Immune phenotyping and batch correction across IMC samples.**

**(a)** Summary table of primary and secondary immune phenotypes, along with functional markers used for profiling.

**(b)** Top: Stacked bar plot showing immune cell fractions across samples from Coh-1 (n = 28) and Coh-2 (n = 14). Bottom: PCA plot showing no clear separation between cohorts. Controls (n = 3) and unsequenced samples (n = 5) are also included.

**(c)** Immune cell phenotyping was performed using lineage and MCL-relevant IMC markers, including CCND1, SOX11, and canonical markers for B, T, and myeloid cells.

**(d)** Top: Uncorrected UMAP showing sample clustering and batch effects. Bottom: UMAP after batch correction with cyCombine shows intermixing of cells from TMA-1, TMA-2, and unstained slides, indicating successful batch correction.

**(e)** Violin plots comparing immune cell fractions across *ATM*-perturbed, *TP53*-perturbed, and WT groups. Group differences were assessed using Wilcoxon rank sum test.

**Supplementary Fig. 10: Development of *in vitro* cell line models for p53 function analysis in MCL.**

(**a**) Ectopic expression of Dox-inducible WT-p53 in Maver-1 and Mino cells. The cells were treated with Dox for 24h, followed by WB examination of p53 and apoptotic markers.

(**b-d**) Establishment of p53-KO or mutations in Z-138 cells using CRISPR technology. Cells were electroporated with various CRISPR constructs followed by GFP sorting and single clone selection. The CRISPR design and representative sequencing confirmation are shown.

**Supplementary Fig.11: Differential Motif Enrichment in Z-138 Isogenic Cells.** The figure shows the top 10 motifs ranked by net target percentage in Z-138 cells with p53-WT, p53-R248Q, and p53-R273C, respectively.

**Supplementary Fig.12: P53-mediated suppression of BCR signaling via upregulation of *PTPN6* (SHP-1).**

**(a)** Assessment of BCR signaling under caspase-inhibited conditions following enforced WT-p53 expression. The pan-caspase inhibitor Z-VAD-FMK (40 μM) was pre-applied to Maver-1 and Mino cells for 1 hour, followed by doxycycline induction (125 ng/mL for Maver-1, 250 ng/mL for Mino) for 24 hours. BCR signaling was examined by WB.

**(b)** Analysis of BCR signaling in Z-138 isogenic cell pairs expressing WT or modified p53. Cells were treated with AMG-232 for 24 h, followed by WB to assess BCR signaling and apoptotic markers.

**(c)** Evaluation of siRNA KD efficiency targeting *PTPN6* in Maver-1 cells. Cells were transfected with various siRNA combinations for 48 hours, followed by WB to confirm KD efficiency.

### Supplementary Fig. 1

**a**

**Coh-1**

| Parameter | OS HR (95% CI) | OS P-Value | PFS HR (95% CI) | PFS P-Value |
|---|---|---|---|---|
| SOX11 IHC | 0.68 (0.33–1.37) | 0.281 | 0.85 (0.42–1.72) | 0.654 |
| TP53 IHC | 2.67 (1.52–4.70) | 0.001 | 1.05 (0.57–1.93) | 0.871 |
| Ki-67 IHC | 1.13 (0.65–1.95) | 0.668 | 1.07 (0.64–1.81) | 0.791 |
| Pathology Subtype | 1.53 (0.70–3.34) | 0.285 | 1.29 (0.60–2.79) | 0.512 |
| Growth Pattern | 1.18 (0.59–2.36) | 0.646 | 1.21 (0.63–2.32) | 0.559 |
| Nodal Status | 0.90 (0.69–1.19) | 0.468 | 1.17 (0.86–1.60) | 0.321 |
| B Symptoms | 0.56 (0.34–0.91) | 0.018 | 0.53 (0.32–0.87) | 0.013 |
| Transplant | 2.46 (1.47–4.12) | 0.001 | 2.00 (1.24–3.22) | 0.005 |
| MIPI | 0.60 (0.20–1.82) | 0.367 | 0.77 (0.26–2.33) | 0.646 |
| Sex | 1.32 (0.78–2.23) | 0.295 | 1.16 (0.72–1.87) | 0.534 |
| Age | 1.57 (1.01–2.45) | 0.047 | 2.05 (1.31–3.22) | 0.002 |
| Rituximab Administered | 0.41 (0.24–0.69) | 0.001 | 0.83 (0.44–1.56) | 0.554 |

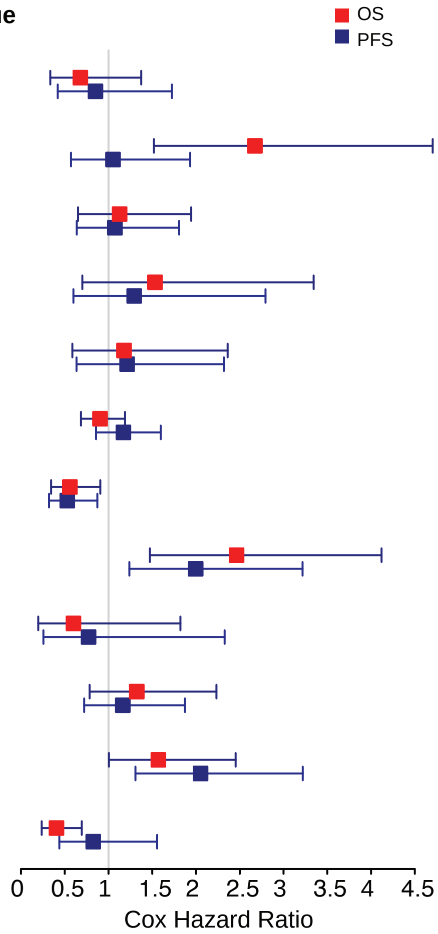


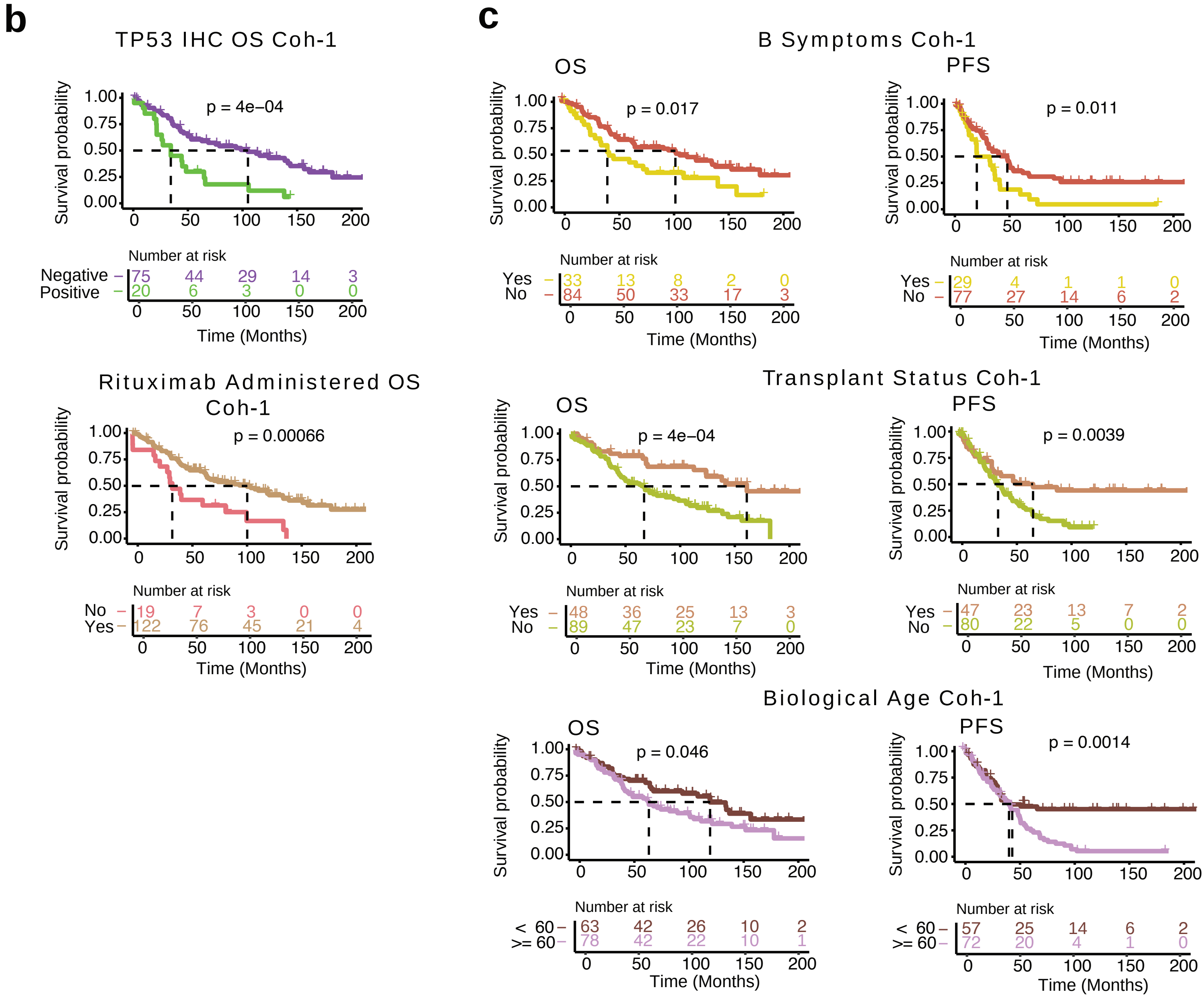

### Supplementary Fig. 2

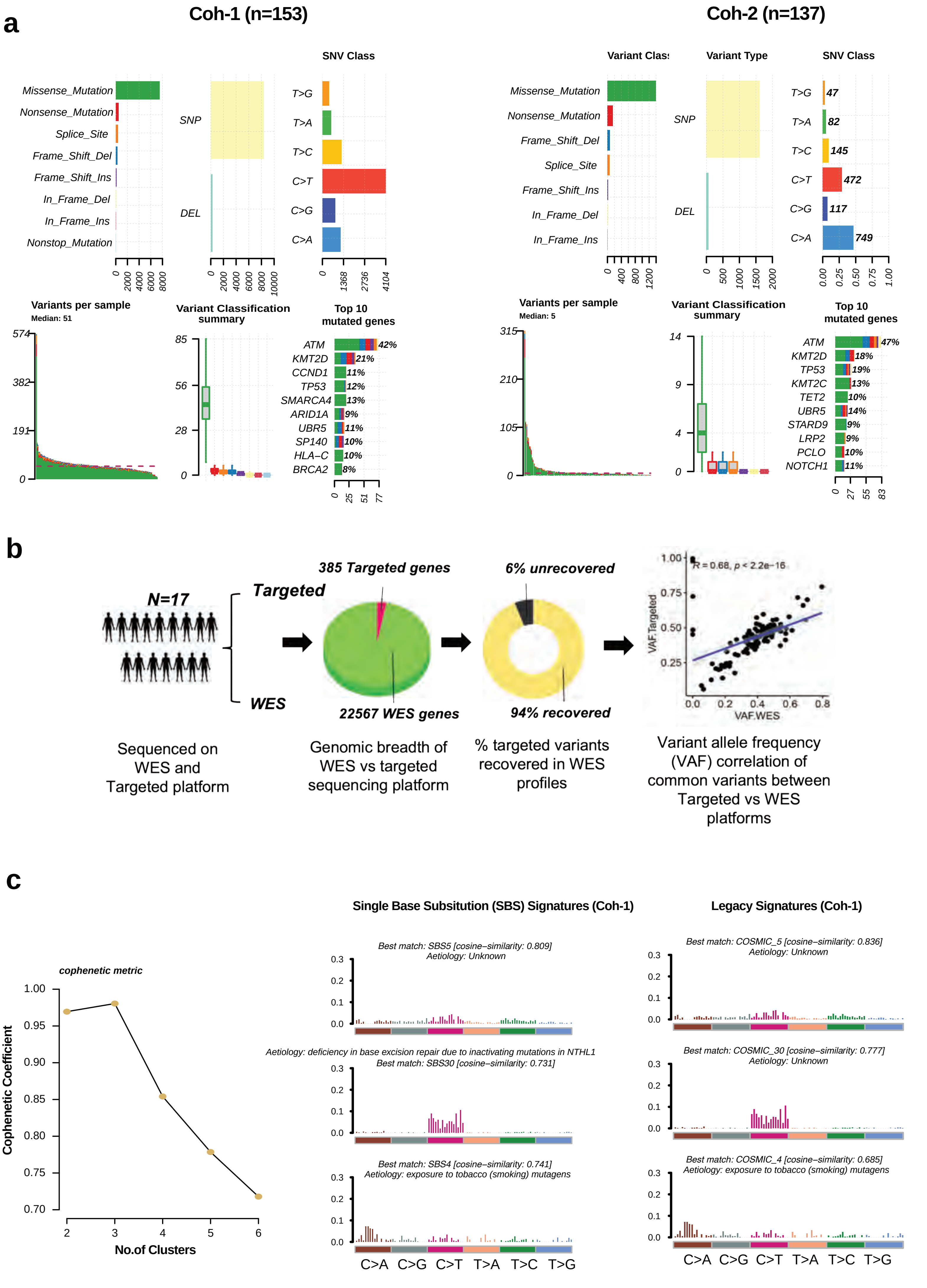

### Supplementary Fig. 3

**a**

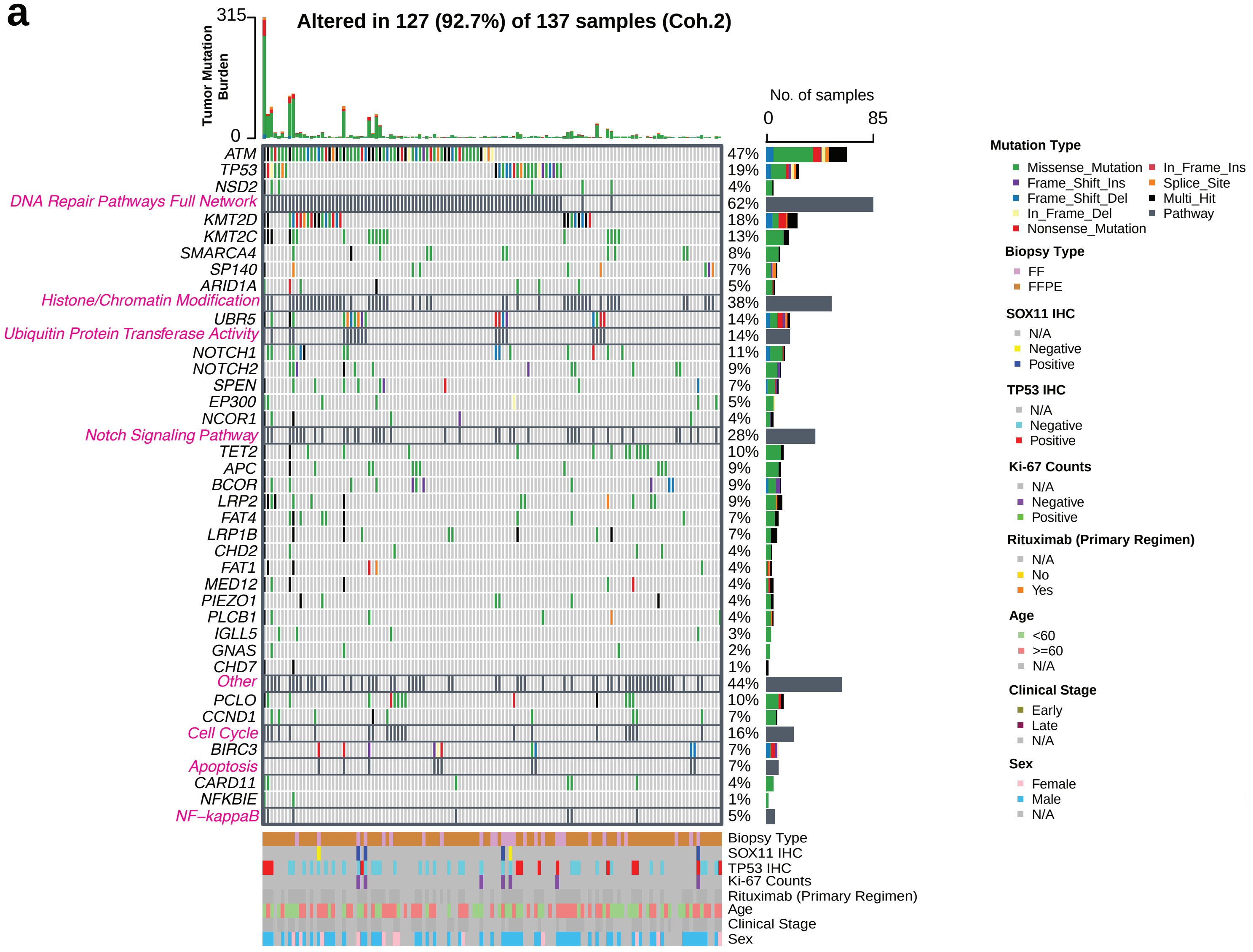


**b**

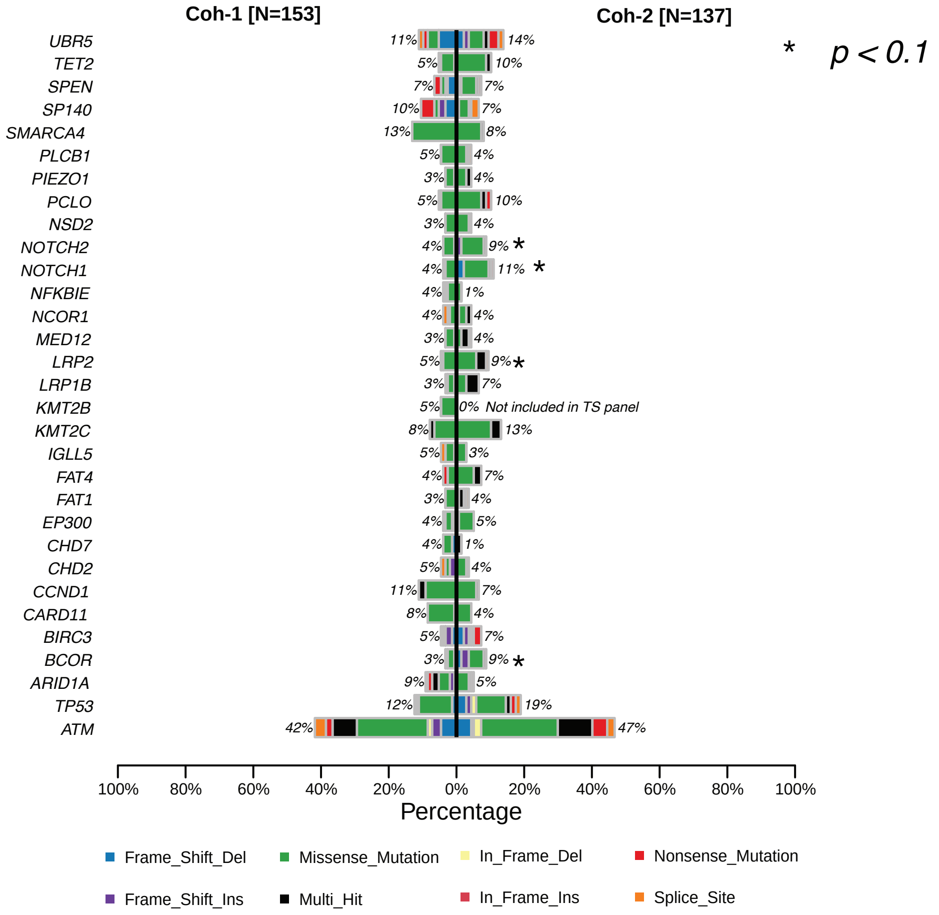

#### Supplementary Fig.4

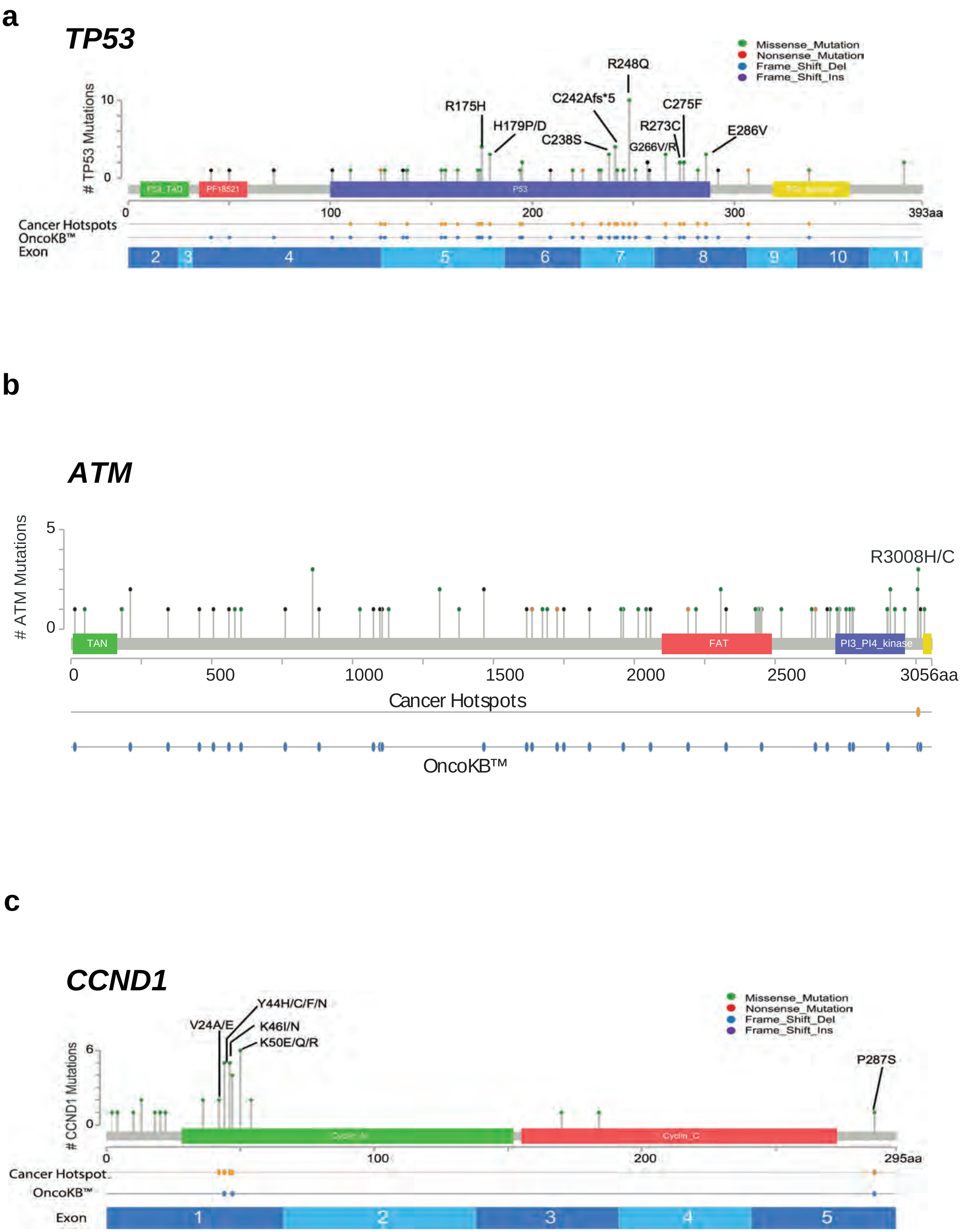

### Supplementary Fig. 5

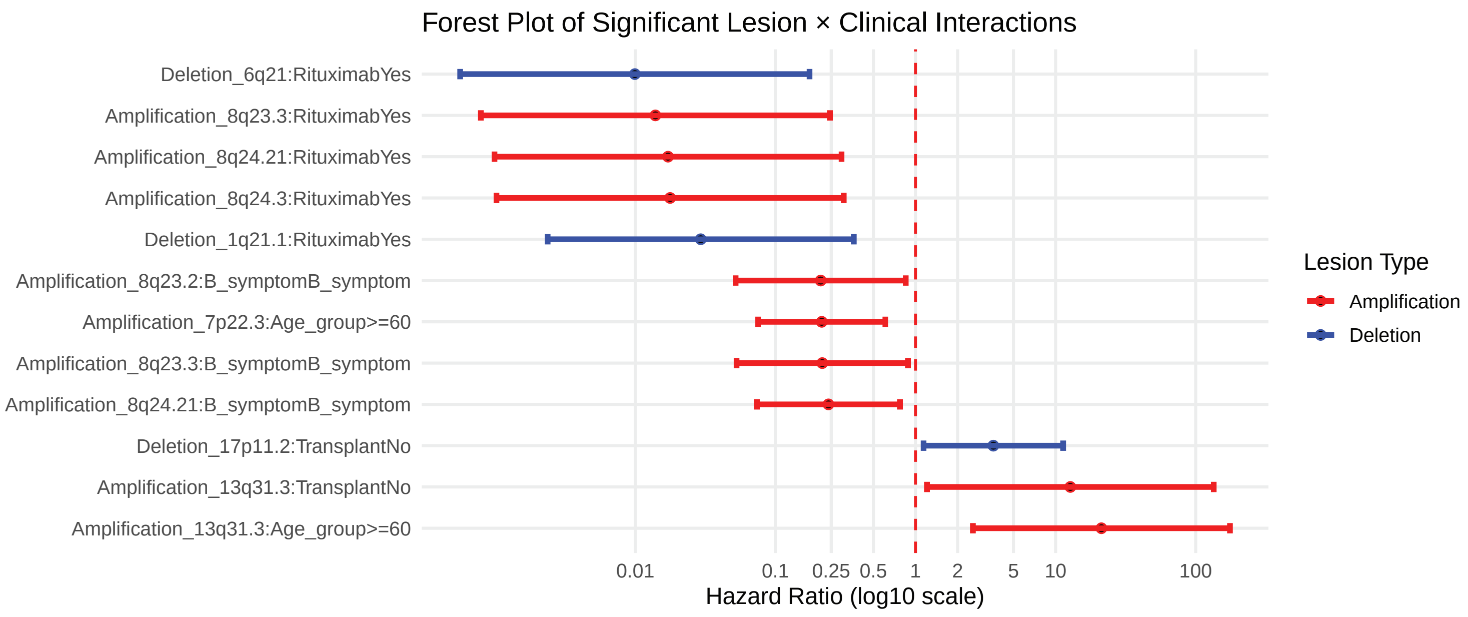

### Supplementary Fig. 6

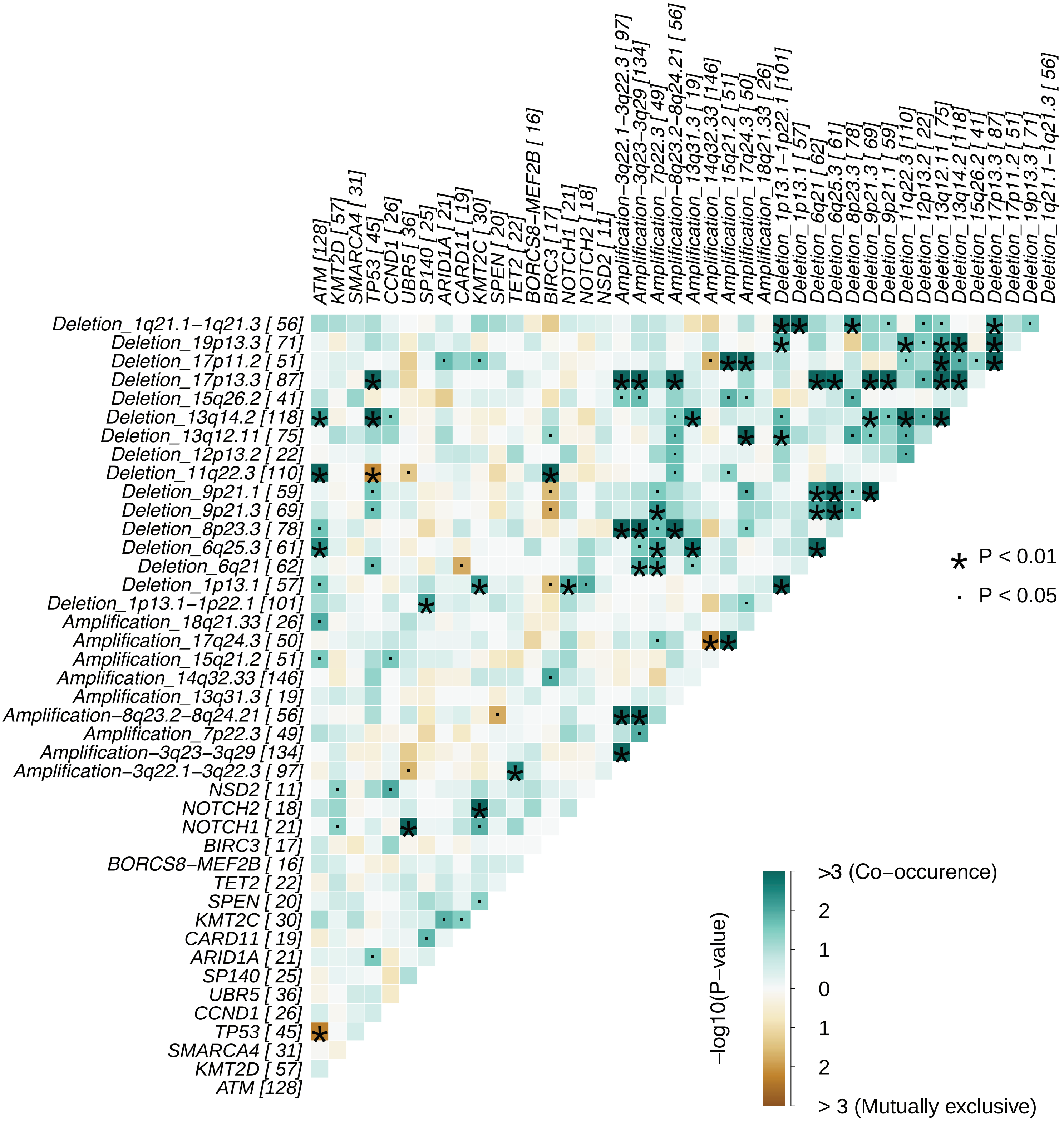

### Supplementary Fig.7

a

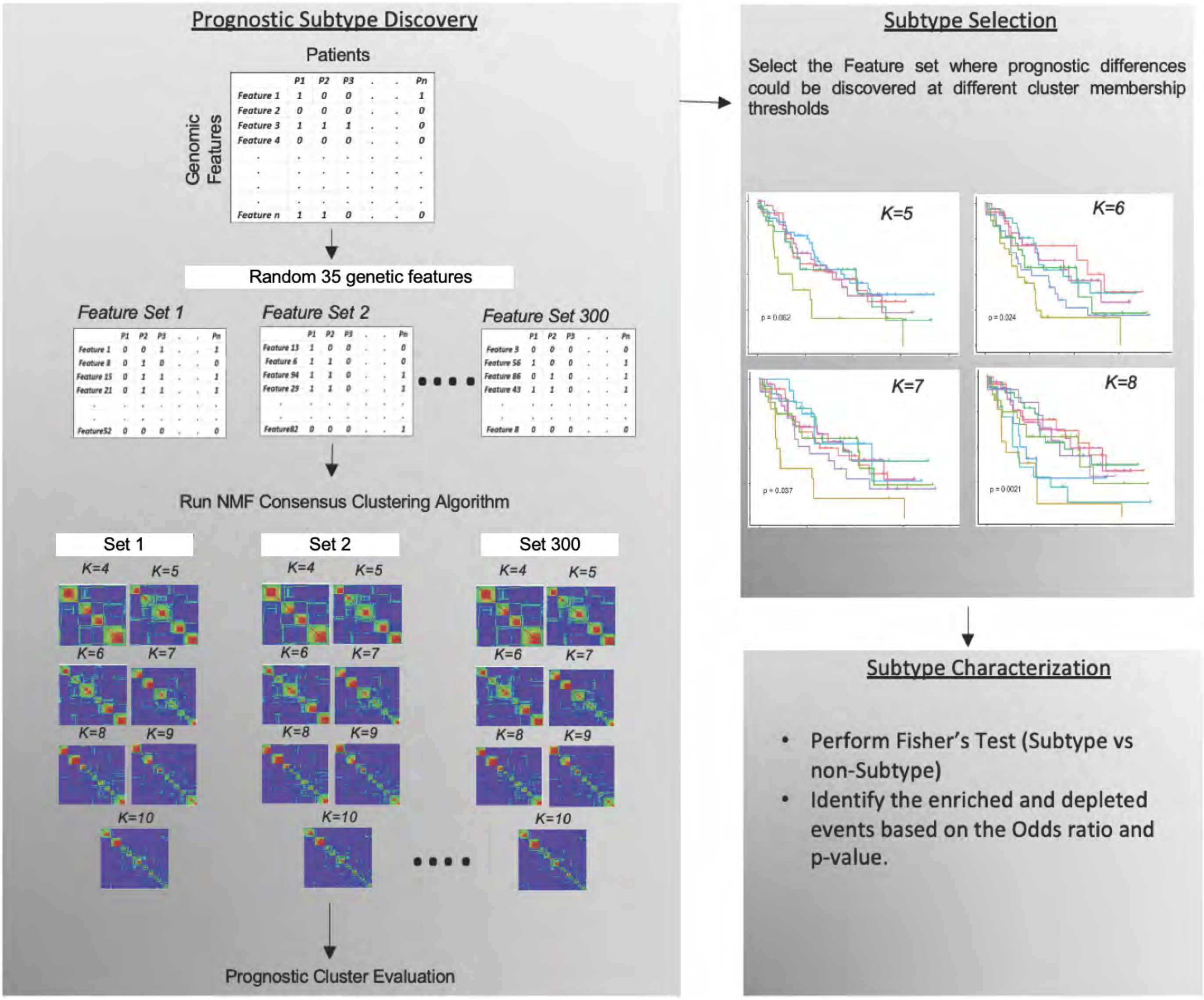


b

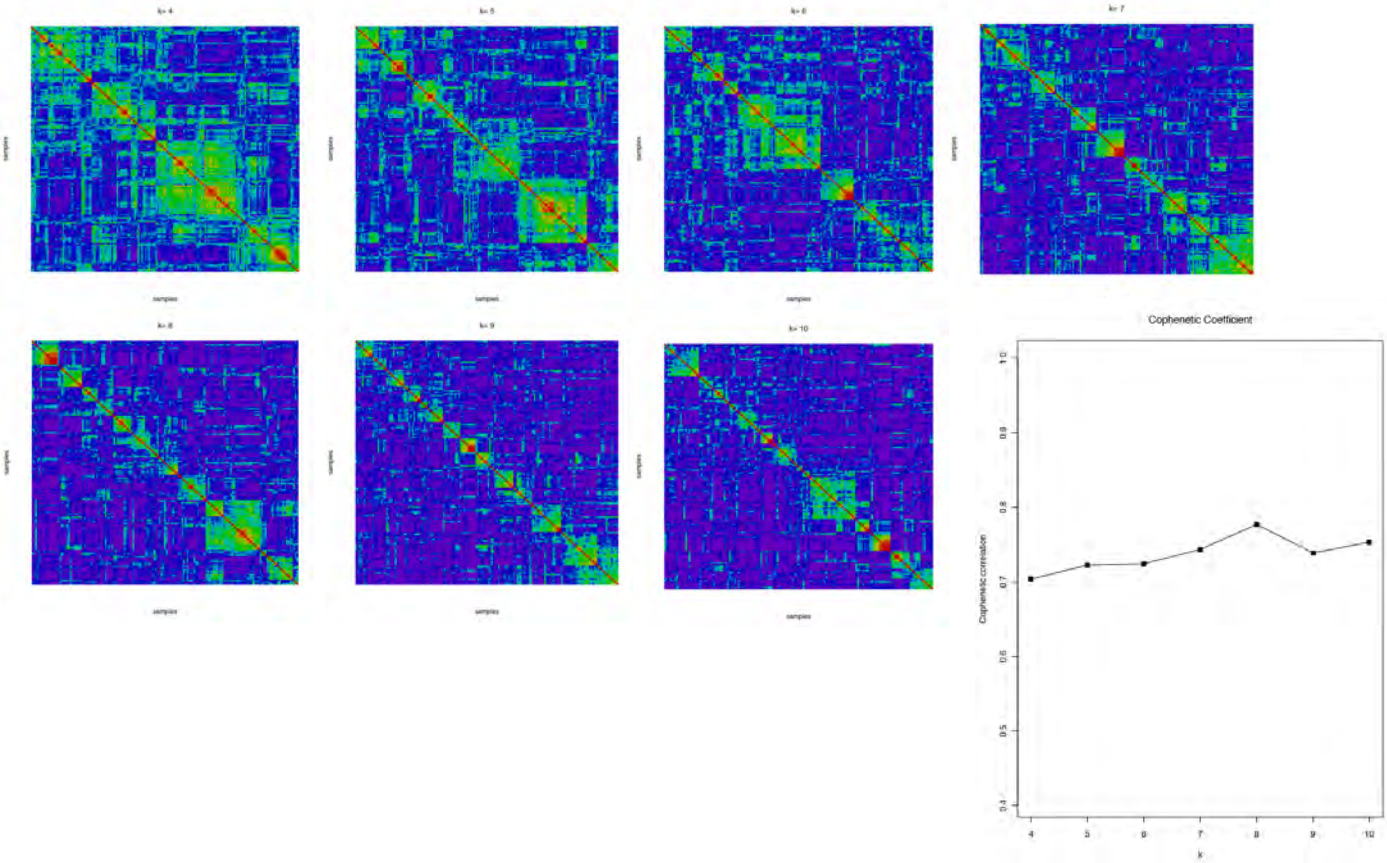

### Supplementary Fig. 8

**a**

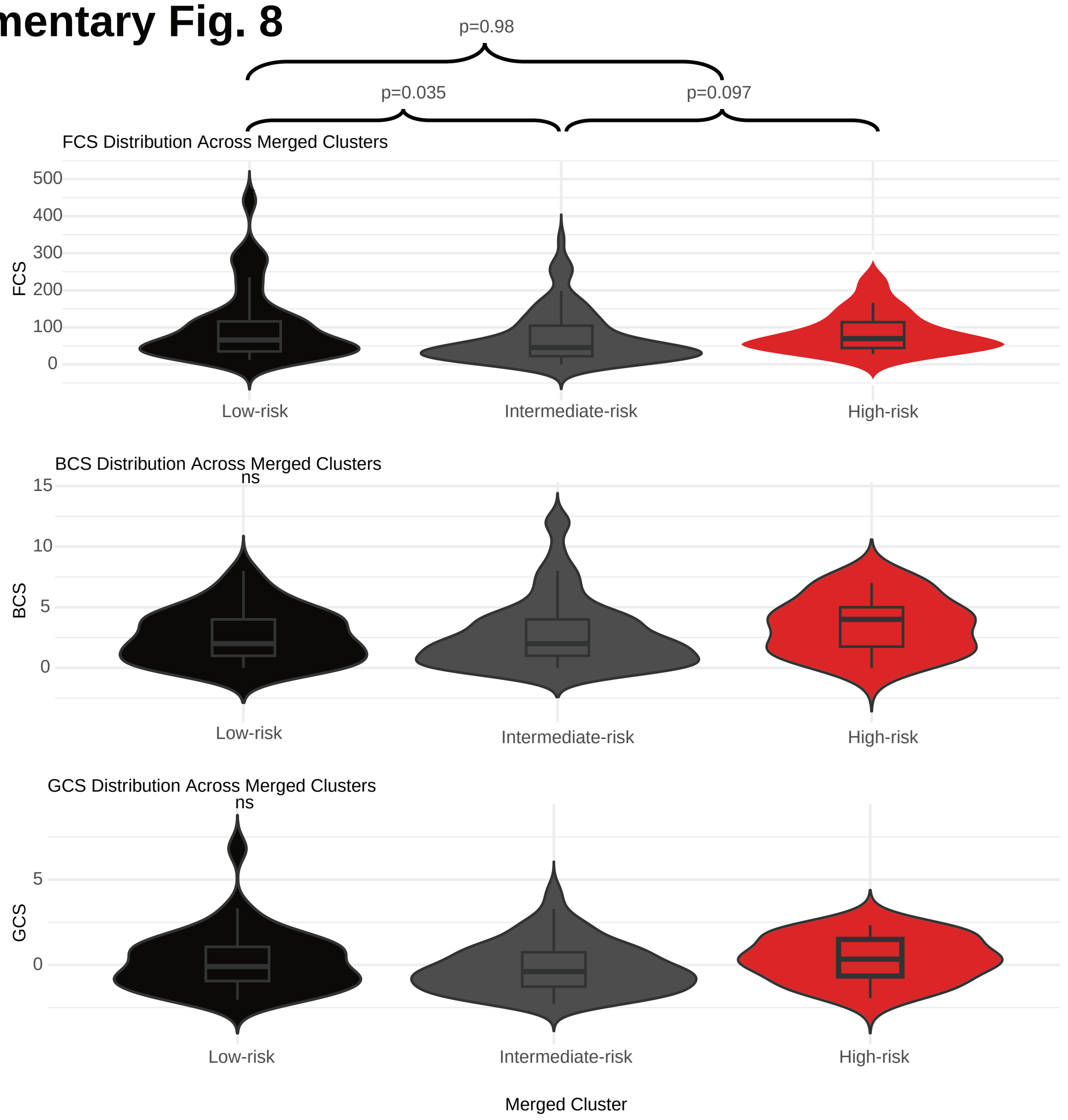


**b**

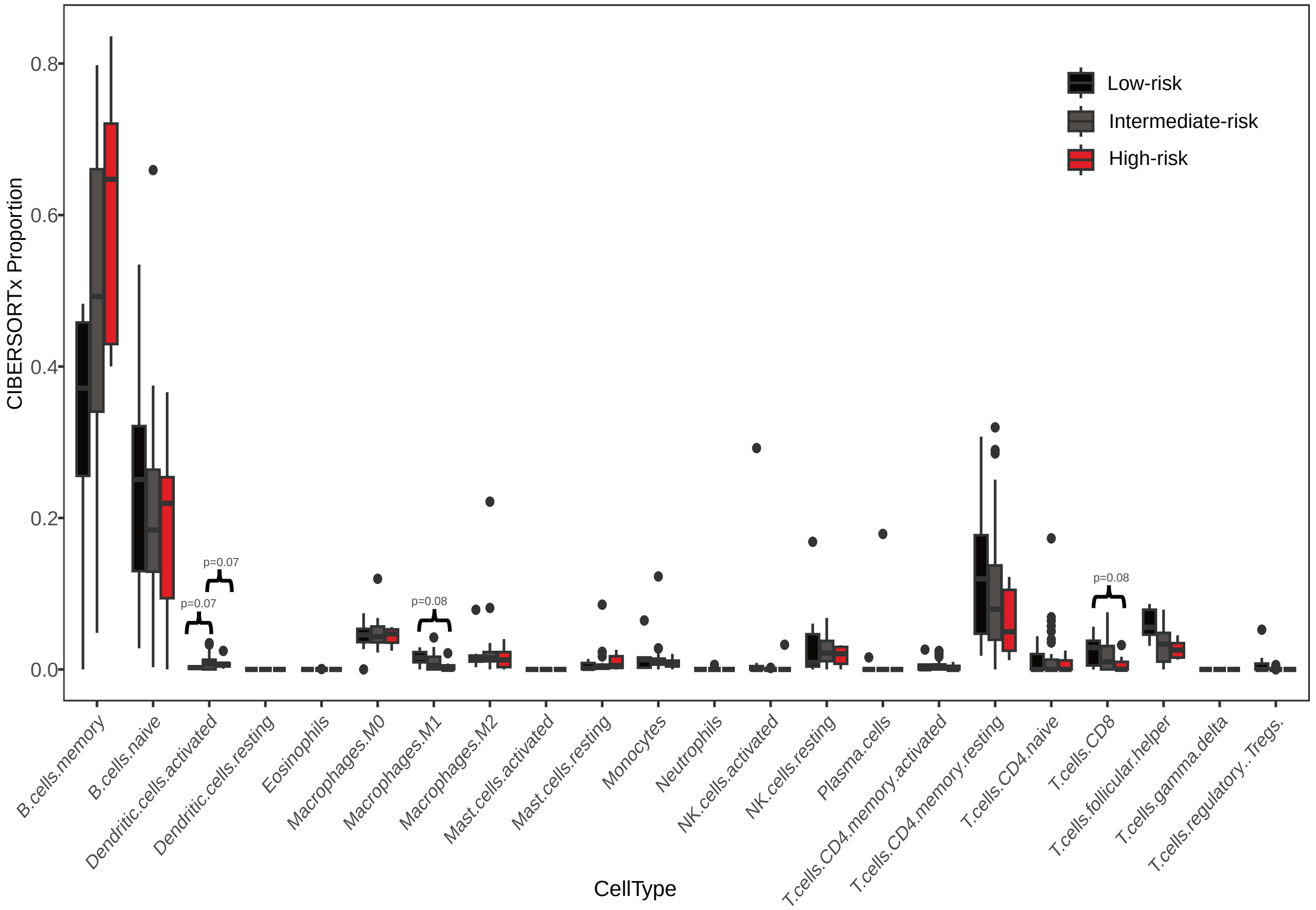

#### Supplementary Fig.9

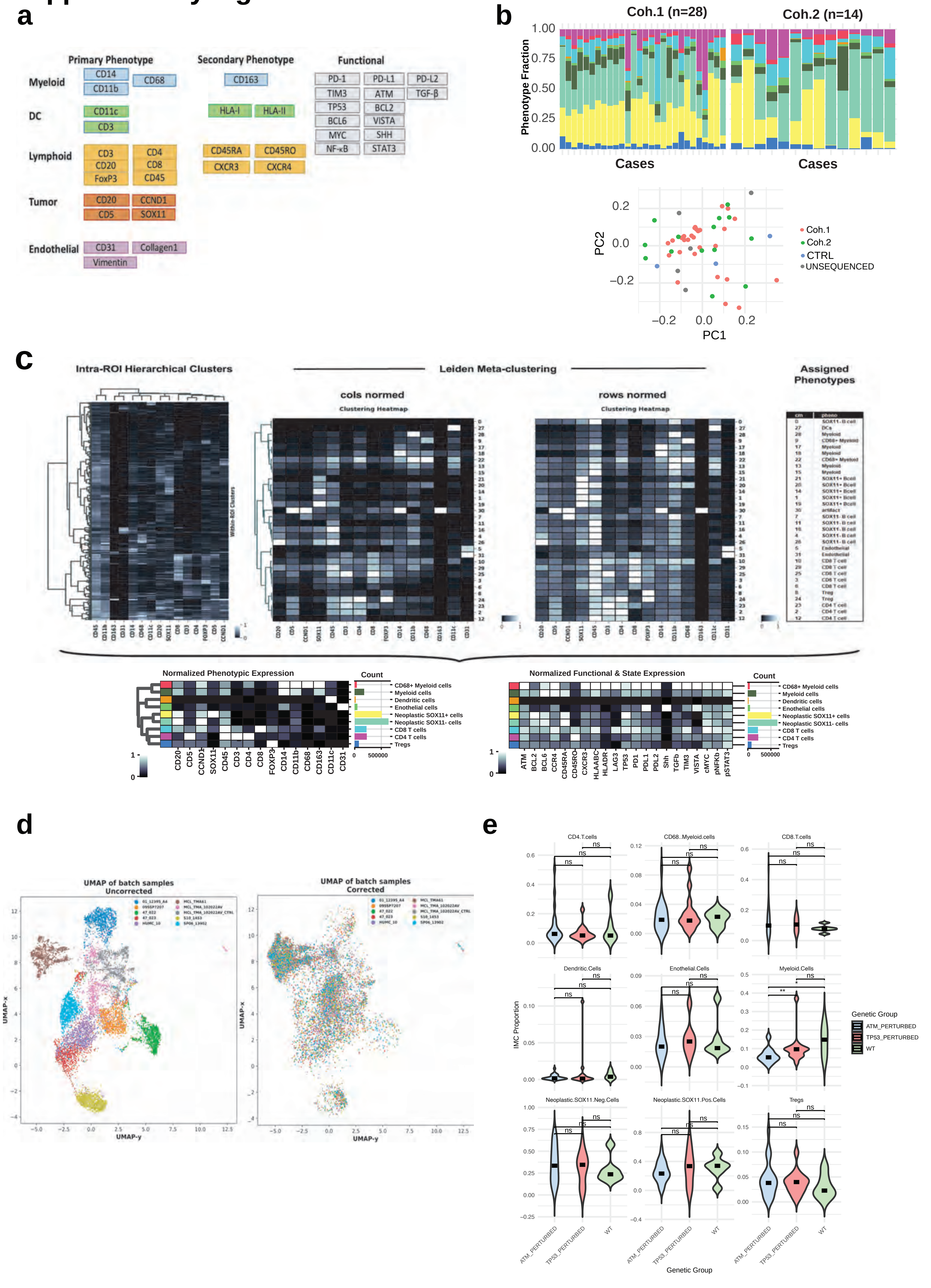

### Supplementary Fig.10

a

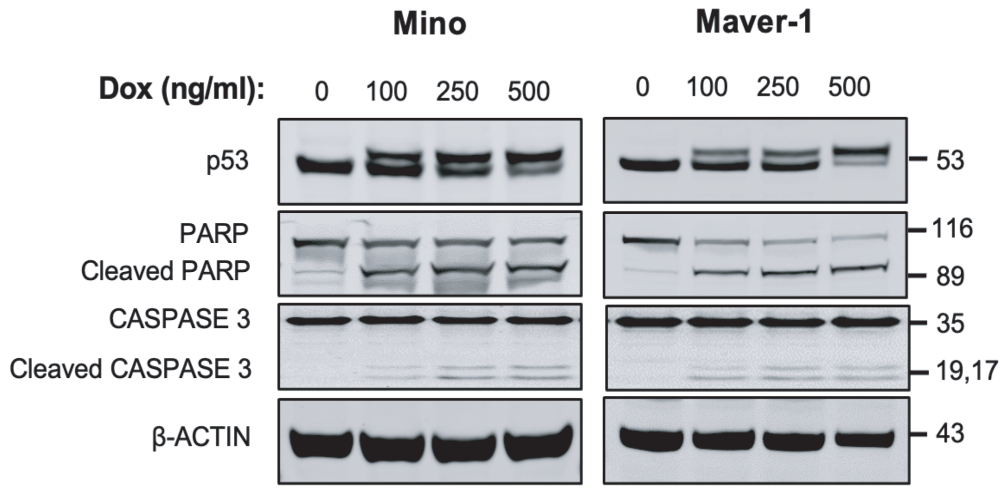


b

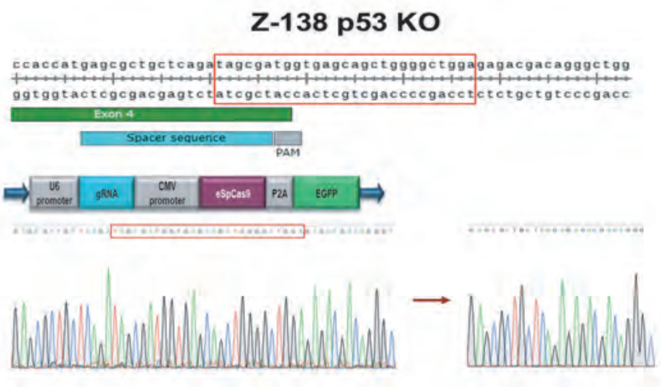


c

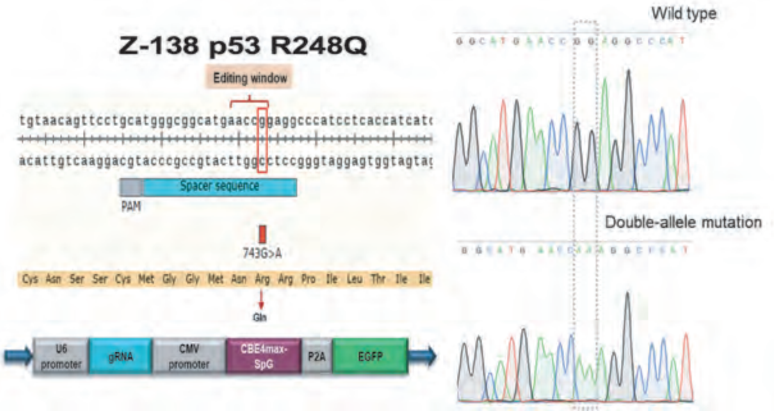


d

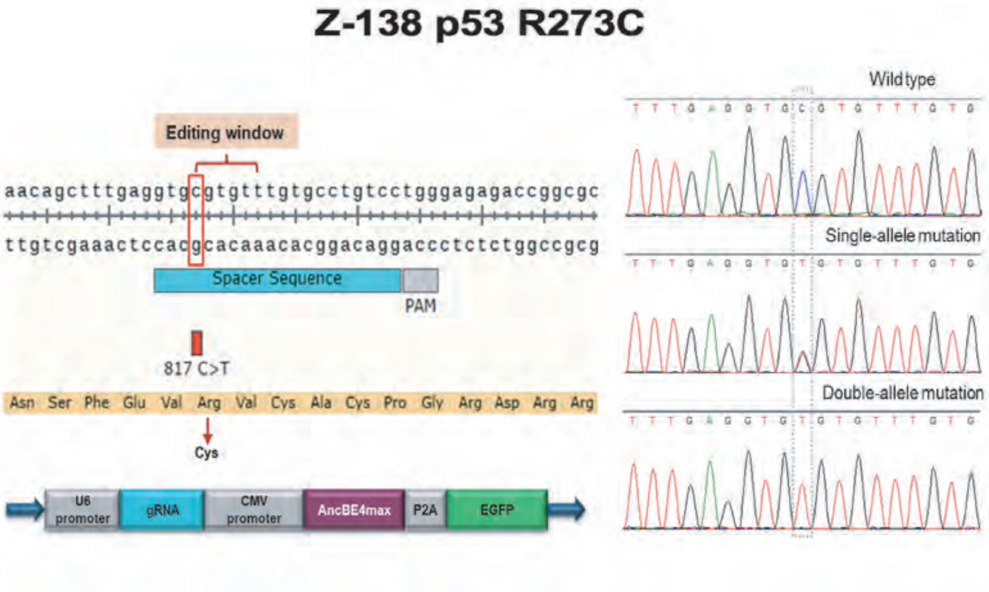

#### Supplementary Fig.11

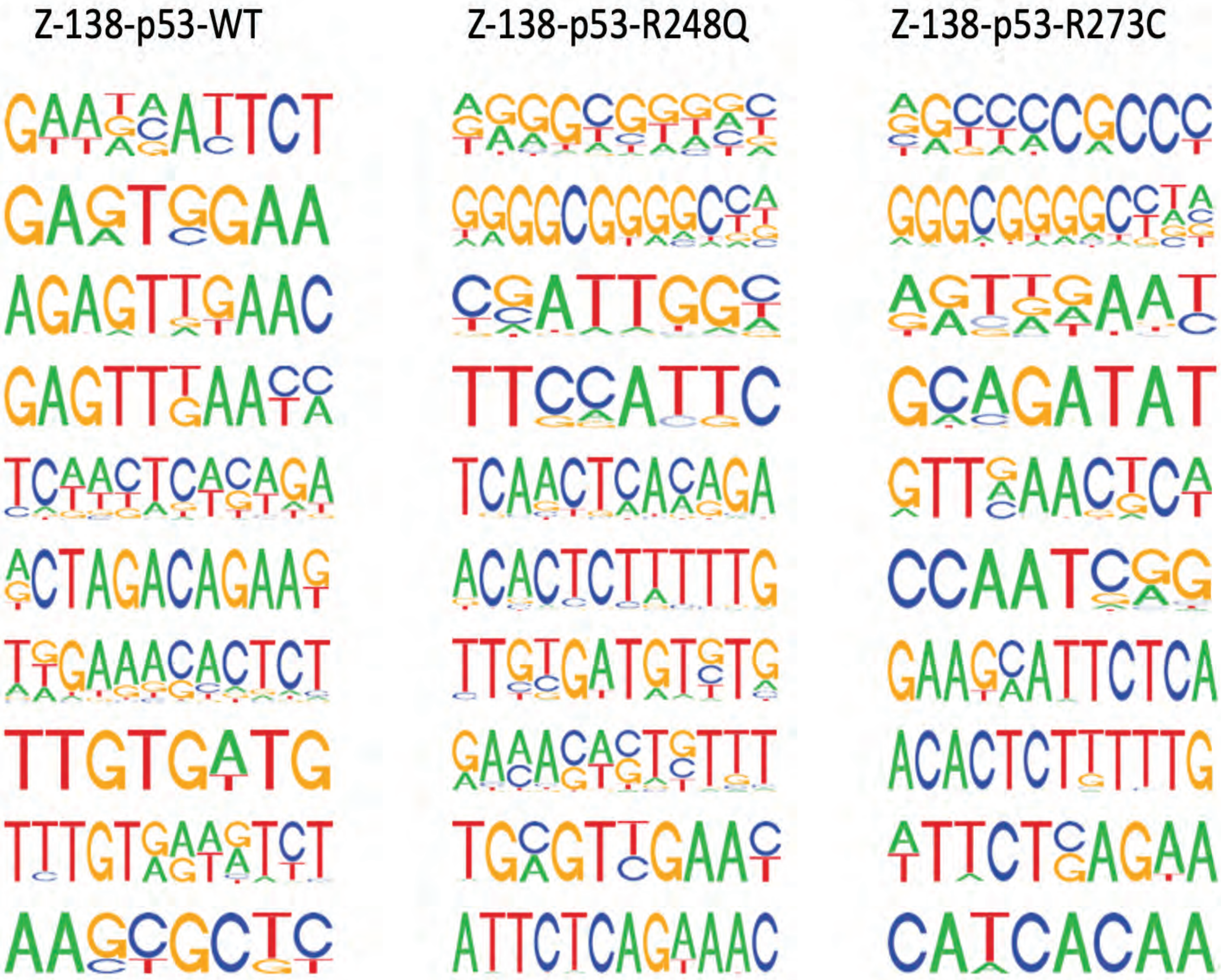

### Supplementary Fig.12

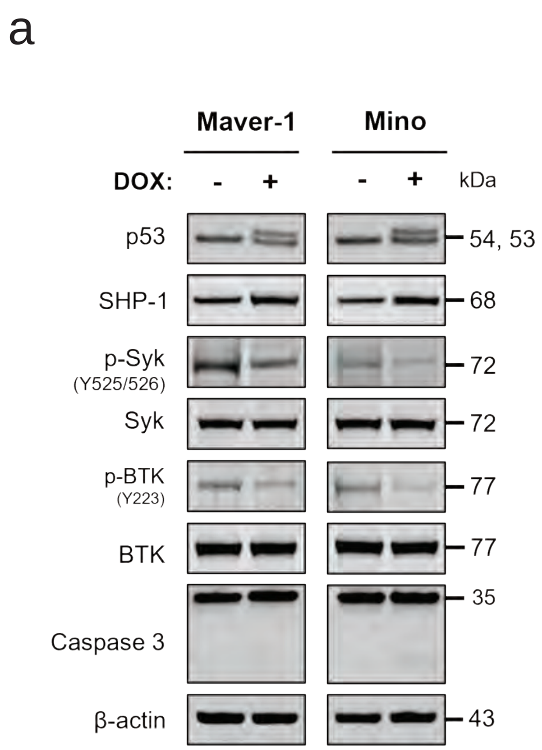


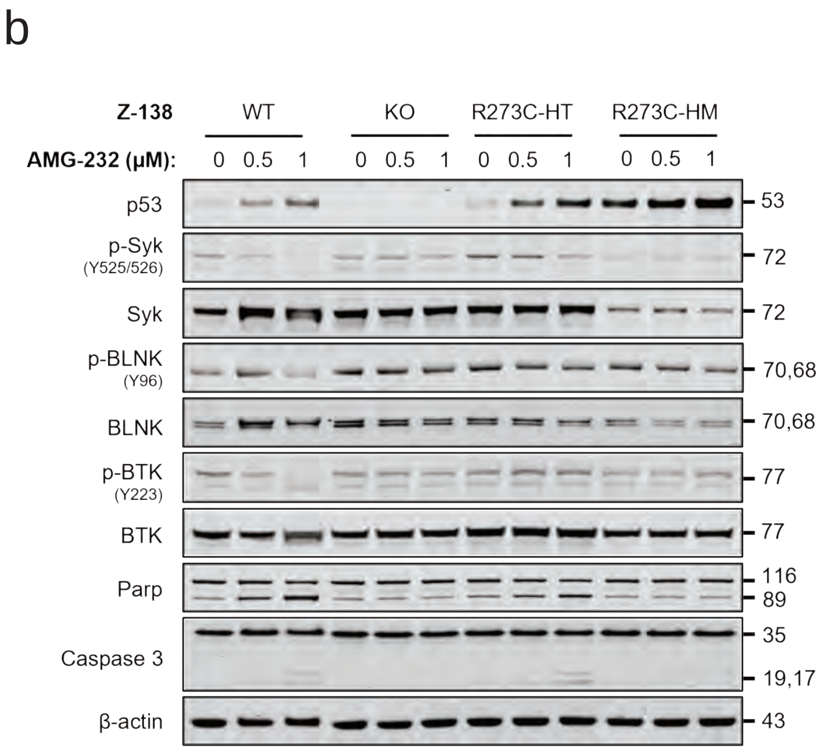


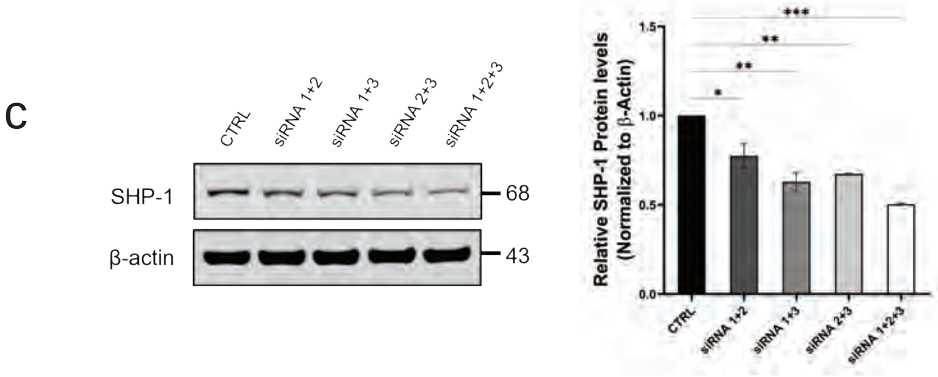
